# Peripheral mutations underlie promiscuous transport of quaternary ammonium antiseptics by Small Multidrug Resistance transporters

**DOI:** 10.1101/2024.02.06.579181

**Authors:** Olive E. Burata, Ever O’Donnell, Jeonghoon Hyun, Rachael M. Lucero, Junius E. Thomas, Ethan M. Gibbs, Isabella Reacher, Nolan A. Carney, Randy B. Stockbridge

## Abstract

The mechanistic basis of transport promiscuity in multidrug exporters is not well understood. We examine this question using the Small Multidrug Resistance (SMR) transporters. We engineer a selective SMR protein to promiscuously export quaternary ammonium antiseptics, similar to multidrug exporters in this family. Using combinatorial mutagenesis and deep sequencing, we identify the necessary and sufficient molecular determinants of this new activity. Using x-ray crystallography, electrophysiology, and a novel proteoliposome-based antiseptic transport assay, we tease apart the mechanistic roles that these residues play in transport polyspecificity. We find that substrate preference changes not through modification of the residues that directly interact with the substrate, but through mutations peripheral to the binding pocket. Our new molecular insights into substrate promiscuity among the SMRs can be applied to understand multidrug export and the evolution of novel transport functions more generally.

## Introduction

The constraints of the two-dimensional membrane environment limit the number of folds available to membrane proteins^1^. And yet, through evolution, a relatively small number of conserved folds have radiated into functionally diverse transporters that are able to translocate all of the structurally and chemically diverse solutes necessary for life^2^. Because large conformational changes are inherent to transporter function, this functional divergence requires alterations to substrate binding in the ground state, as well as the entire structural and energetic landscape during the conformational change. Understanding the molecular basis of how novel transport functions emerge through evolution is thus a challenging question.

We have established proteins from the Small Multidrug Resistance (SMR) family as a tractable system to analyze how changes in substrate specificity arise through molecular events. These proteins are among the smallest characterized membrane transport proteins, with only one hundred residues total. Pairs of identical subunits assemble as antiparallel homodimers^3^. This assembly gives rise to a symmetric energy landscape with structurally and energetically equivalent inward- and outward-facing structures^4^. This primitive architecture was a likely evolutionary antecedent to the pseudosymmetric inverted repeat architecture that is common among secondary active transporters^5^. Thus, the SMRs provide a general model for early evolution of transport proteins^6^.

SMR transporters are widespread among microorganisms, and have evolved several distinct functional subtypes^7^. Two of these are considered in this manuscript: the SMR_Gdx_ (**<u>g</u>**uani**<u>d</u>**inium e**<u>x</u>**port) and the SMR_Qac_ (**<u>q</u>**uaternary **<u>a</u>**mmonium **<u>c</u>**ation). The SMR_Gdx_ selectively transport guanidinium ion (Gdm^+^), a small cationic byproduct of nitrogen metabolism^8^, along with close polar analogs like guanylurea^9,10^. In contrast, transporters from the SMR_Qac_ subtype are promiscuous exporters of hydrophobic cationic compounds^11–13^, including quaternary ammonium compounds like benzalkonium and cetyltrimethylammonium (also known as cetrimonium or CTA^+^), the main antiseptic agents in common household and hospital cleaning solutions and antibacterial handsoaps^14^. Their genes are among the most frequent associated with horizontal transfer of drug resistance among bacterial populations^15–17^. The SMR_Qac_ subtype includes the well-studied multidrug exporter from *Escherichia coli*, EmrE. Despite their substantial differences in substrate transport specificity, transporters from the SMR_Gdx_ and SMR_Qac_ subtypes exhibit high structural (1.2 Å C_α_ RMSD) and sequence (∼40%) similarity^7,18^. The proteins share an absolutely conserved pair of central glutamates at the bottom of the substrate binding pocket, which bind the cationic substrate or protons in a mutually exclusive fashion^19^. In fact, most of the residues that line the substrate binding pocket are conserved, raising the question of how these substantial changes in substrate selectivity arose without altering the protein residues in closest proximity to the substrate.

Comparative structural studies of EmrE and the best-studied SMR_Gdx_, from *Clostridales* oral taxon 876, called Gdx-Clo, revealed key differences in the interactions between these conserved residues^10,18^. Whereas the binding pocket sidechains in Gdx-Clo are organized in a polarized hydrogen bond network that stabilizes the substrate- and proton-binding central glutamates, in EmrE, a weak and disorganized hydrogen bond network yields more rotameric freedom for binding pocket sidechains^18^. EmrE structures solved in complex with substrates of varying bulk and aromatic character show that this flexibility – especially of the central glutamates (E13) and binding site aromatic W63 – accommodates structurally dissimilar substrates^18^. Despite these recent structural advances, the molecular determinants of promiscuous substrate transport in EmrE remain unknown. Numerous mutant proteins have been examined, and perturbations to transport selectivity have been described for many of them^20–22^. At the same time, a series of biophysical experiments have established detailed information about EmrE’s dynamic properties^4,23–27^, which have been invoked to explain transport promiscuity. However, these studies have not coalesced into a unified understanding of the specific protein sequences that underlie the emergence of promiscuous transport behavior characteristic of the SMR_Qacs_ within the larger SMR family.

In order to understand the molecular features responsible for substrate polyspecificity among SMRs, we engineer a Gdx-Clo variant that is capable of transporting structurally dissimilar quaternary ammonium antiseptics. Using combinatorial mutagenesis, we establish a set of residues that are necessary and sufficient for this activity. These residues are conserved among SMR_Qacs_, supporting the interpretation that these residues also reflect natural mutations that led to promiscuous transport by the SMR_Qacs_. By combining structural studies, binding, transport, and electrophysiology assays, we establish the mechanistic basis for quaternary ammonium antiseptic transport by this family of transporters.

## Results

### An engineered variant of Gdx-Clo confers resistance to quaternary ammonium antiseptics

Analysis of sequence alignments of the SMR_Qac_ and SMR_Gdx_ transporters^7^ revealed six residues that are highly conserved in each subtype, but that differ between the subtypes. These residues were, for Gdx-Clo and EmrE, respectively, Gly10/Ile11, Trp16/Gly17, Ala17/Thr18, Met39/Tyr40, Ala67/Ile68, and Lys101/Asn102. We reasoned that these residues might be mainly responsible for the differing substrate selectivity of the SMR_Qac_ and SMR_Gdx_ subtypes. We therefore introduced the SMR_Qac_ residues at all six positions in Gdx-Clo (G10I, W16G, A17T, M39Y, A67I, and K101N), but the resulting transporter failed to confer bacterial resistance to quaternary ammonium antiseptics.

To identify additional mutations that might furnish quaternary ammonium resistance and substrate polyselectivity, we further subjected this mutated Gdx-Clo construct to random mutagenesis followed by selection for bacterial growth in the presence of two different quaternary ammoniums, cetrimonium (CTA^+^) and tetrapropylammonium (TPA^+^) to select for promiscuous transporters. We chose these substrates because they differ in the bulkiness of the quaternary ammonium group, and cetrimonium (but not TPA^+^) possesses a long alkyl tail characteristic of many common quaternary ammonium antiseptics. Moreover, the differing inhibitory concentrations of these two compounds (∼200 μM and ∼20 mM, respectively) demand export efficiency over ∼2 orders of magnitude concentration. Under these selection conditions, one round of directed evolution yielded a construct that supported robust growth of Δ*emrE E. coli* in the presence of both CTA^+^ and TPA^+^. This construct bore the six original rationally designed mutations and an additional seventh mutation, A60T. The resistance conferred by this construct, dubbed Gdx-Clo-7x, is comparable to that of native EmrE introduced on a rescue plasmid (**Figure 1A**). By themselves, none of the seven mutants, including A60T, exhibit quaternary ammonium resistance (**Supplementary Figure 2**).

**Figure 1.**
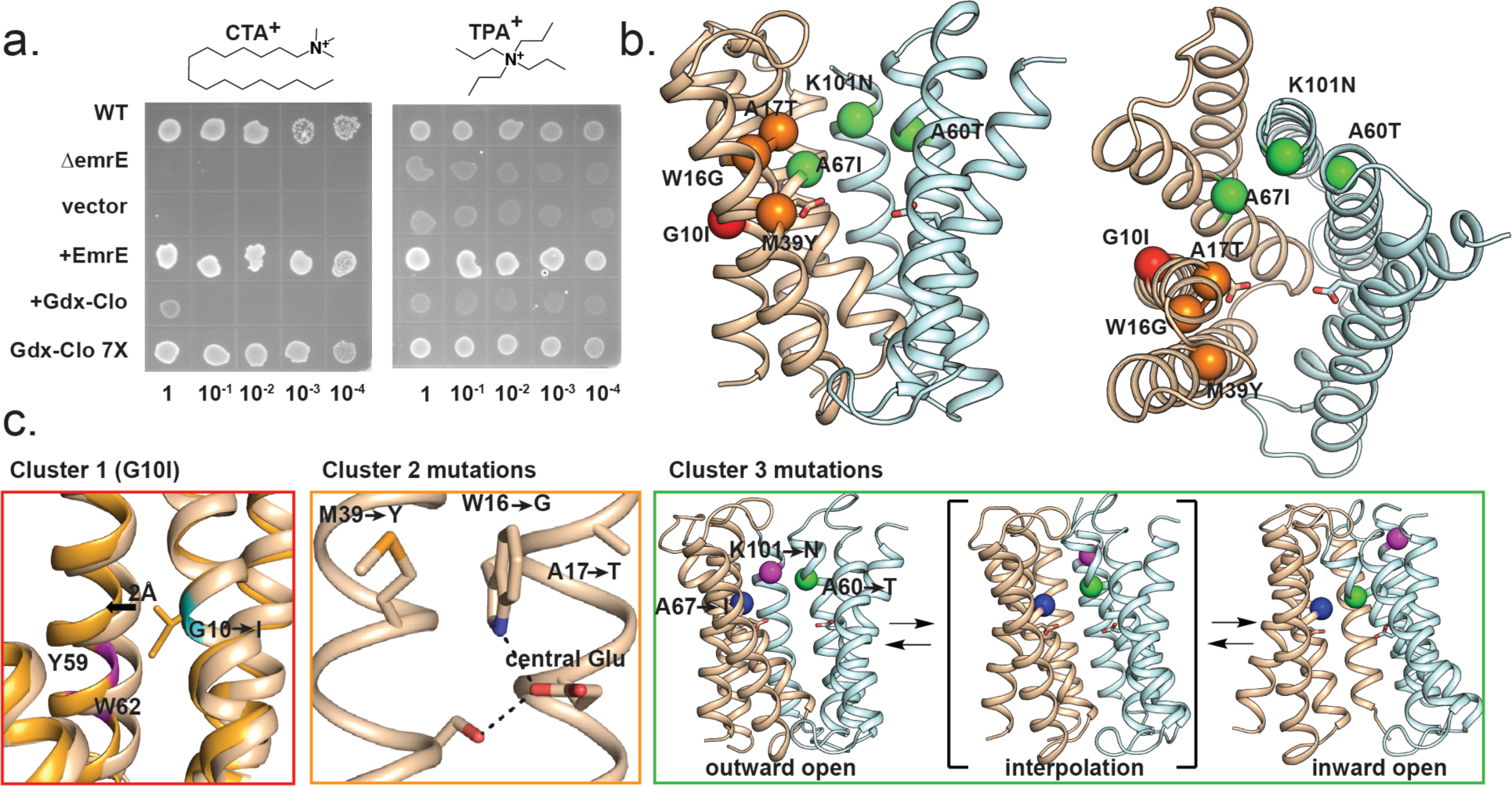
An engineered variant of Gdx-Clo confers bacterial resistance to quaternary ammonium antiseptics. A) Serial dilutions of WT (top row) or Δ*emrE E. coli* (bottom 5 rows) bearing SMR variants as indicated. Selections were performed with 120 µM CTA^+^ (left) or 18 mM TPA^+^ (right). Control dilutions on no-drug control plates are shown in **Supplementary Figure 1**. B) Structure of WT Gdx-Clo (PDB: 6WK8) with C_α_ atoms of the Gdx-Clo-7x mutations shown as spheres. The A and B monomers are show in pale cyan and tan, respectively, and the central glutamates are shown as sticks. For clarity, mutations are shown on only one monomer. Mutations are colored according to structurally co-localized cluster, as described in the text (cluster 1: red; cluster 2: orange; cluster 3: green). C) Left, overlay of the B monomer of Gdx-Clo (tan; PDB:6WK8) and EmrE (orange; PDB: 7MH6) with the position of cluster 1 mutation (G10/I11) shown in cyan. The positions of binding site residues Y59 and W62 (Gdx-Clo numbering) are shown in magenta. Middle: WT Gdx-Clo with positions of cluster 2 mutations, the central glutamate, and S42 shown as sticks. Hydrogen bond interactions are shown as dashed lines. Right: Interpolation of outward- and inward-facing Gdx-Clo with C_α_ atoms of cluster 3 mutations shown as spheres (A67 in blue, K101 in magenta, and A60 in green).

The seven mutations of Gdx-Clo-7x identified are distributed across the protein in three structurally co-localized clusters (**Figure 1B**). The first, G10I (red), occurs at a helical packing motif between helices 1 and 3 (**Figure 1C**, **left**). These helices contribute several residues to the binding site hydrogen bond network, suggesting that the G10I mutation alters the packing of these two helices and disrupts hydrogen interactions among substrate binding site sidechains. We previously proposed that disruption of the hydrogen bond network in the binding site permits the binding of structurally diverse substrates^18^. The second mutant cluster is comprised of W16G, A17T, and M39Y (orange), located adjacent to the substrate binding site (**Figure 1C**, **center**). W16 is the only one of the seven mutant residues that participates in the hydrogen bond network of WT Gdx-Clo’s substrate binding site, where it directly coordinates the central glutamate, E13. In WT Gdx-Clo, hydrophobic sidechains A17 and M39 sandwich W16 in place. A60T, A67I, and K101N (green) comprise the third cluster, which is located at the periphery of the aqueous binding pocket (**Figure 1C**, **right**). These residues undergo large changes in position during the inward-to outward-facing conformational change, and interpolation of the inward- and outward-facing structures suggests that they might pass in close proximity during the conformational transition.

### Gdx-Clo-7x binds and transports quaternary ammonium substrates, but not Gdm^+^

We next examined the functional and biochemical properties of Gdx-Clo-7x. Size exclusion chromatography showed a monodisperse peak with protein yield comparable to WT Gdx-Clo (**Supplementary Figure 3**). We first assessed substrate binding by monitoring substrate-dependent changes to the proteins’ intrinsic tryptophan fluorescence^9,28^ (**Figure 2A-C**). The seven mutations induce a complete switch in substrate preference for Gdm^+^ and TPA^+^: Gdx-Clo WT binds Gdm^+^ with a K_d_ of 800 µM, but not TPA^+^. In contrast, Gdx-Clo-7x binds TPA^+^ with a K_d_ of 15 mM, in line with the inhibitory concentration used for the bacterial selection experiments, but not Gdm^+^ (K_d_ >50 mM). Both WT and mutant transporter bind CTA^+^ with comparable affinities (7.2 µM for WT Gdx-Clo and 4.7 µM for Gdx-Clo-7x), in agreement with previous data showing detergent binding by members of the SMR_Gdx_ subtype^29^. Together, these binding experiments confirm Gdx-Clo-7x’s interaction with both quaternary ammonium substrates, and show that the expanded recognition of quaternary ammonium compounds is accompanied by greatly diminished binding to Gdm^+^.

**Figure 2.**
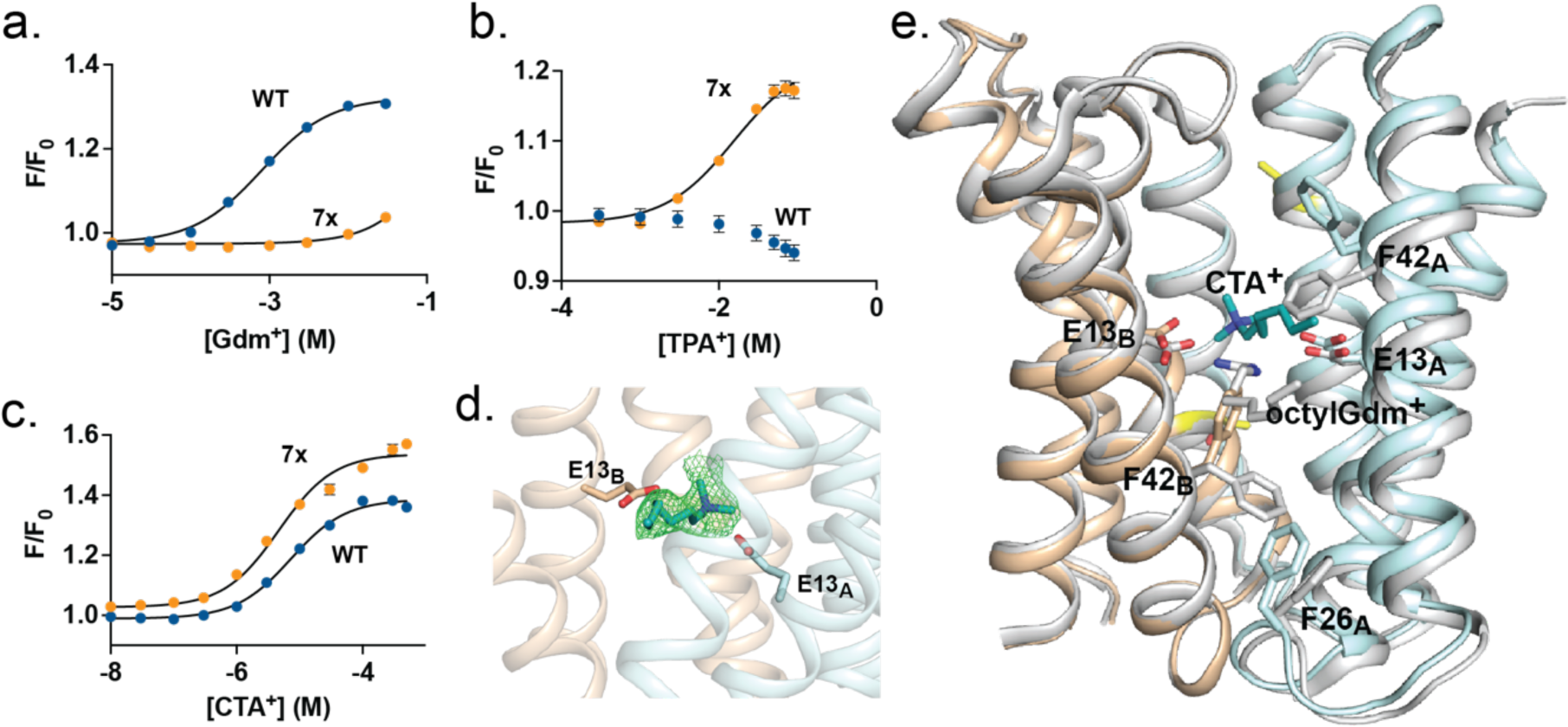
Substrate binding to WT Gdx-Clo and Gdx-Clo-7x. Tryptophan fluorescence as a function of (A) Gdm^+^, (B) TPA^+^, or (C) CTA^+^ concentration, normalized relative to fluorescence intensity in the absence of substrate, for WT Gdx-Clo (blue) and Gdx-Clo-7x (orange). Each datapoint represents the mean and SEM of three independent experiments (where not visible, error bars are smaller than the diameter of the point). Solid lines represent fits to a single-site binding model. K_d_ values derived from these fits are reported in the text. Raw fluorescence spectra are shown in **Supplementary Figure 4**. Data from an independent protein preparation are shown in **Supplementary Figure 5**. D) Structure of CTA^+^ bound to Gdx-Clo A60T. A and B subunits are shown in pale cyan and tan, respectively, and the central glutamates are shown as sticks. Modelled atoms of CTA^+^ are shown in teal, with the F_o_-F_c_ polder omit map, contoured at 4σ, in green. E) Overlay of Gdx-Clo A60T with CTA^+^ bound (colored as in panel D) and WT Gdx-Clo with octylGdm^+^ bound (PDB:6WK9; shown in light gray). The A60T residues are colored in yellow.

We sought to examine the structural basis for quaternary ammonium binding, but we were unable to produce crystals that diffracted to high resolution for Gdx-Clo-7x, or for WT Gdx-Clo in the presence of CTA^+^. We screened some of the single mutant constructs for crystallization and diffraction, and obtained serviceable crystals for Gdx-Clo A60T that allowed us to determine its structure in complex with CTA^+^ (**Supplementary Table 1**). The overall structure was nearly identical to WT Gdx-Clo (C_α_ RMSD = 0.24 Å), and the positions and interactions among major binding site residues (E13, W16, Y59, W62) are unchanged relative to WT Gdx-Clo. We observe only minor perturbations to the structure in the vicinity of the A60T residues, which are located ∼10 Å from, and face away from, the binding site. Moreover, functional experiments show that A60T transports Gdm^+^ at levels similar to WT Gdx-Clo (**Supplementary Figure 6**). We therefore interpret this structure to provide insight into how WT Gdx-Clo binds CTA^+^, as seen in Figure 2C. We observed an elongated density in the binding pocket, into which we modelled the quaternary ammonium headgroup and first seven carbons of the alkyl tail of CTA^+^ (**Figure 2D**). The quaternary ammonium headgroup is located in the vicinity of the central glutamates, but binds about 2.5 Å higher, and about 4 Å farther back in the pocket than do the transported guanidinyl substrates, which are poised in the same plane as the glutamates^10^ (**Figure 2E**). The alkyl tail of CTA^+^ extends towards the membrane, exiting the binding site through a gap between helices 2_A_ and 2_B_, in the manner observed for the extended alkyl tail of octylGdm^+10^. Due to the slightly higher position of the substrate within the binding pocket, the hydrophobic residues that line the membrane portal, notably F42_A_ and F42_B_, rearrange to permit the tail to access the membrane (**Figure 2E**). This structure suggests that although some quaternary ammoniums with small headgroups can be accommodated within the WT binding pocket, these substrates are unable to access its deepest point between the central glutamates, which could plausibly impair their ability to induce conformational change of the transporter.

We further assessed transport of TPA^+^ by Gdx-Clo-7x using solid-supported membrane (SSM) electrophysiology (**Figure 3A**). Due to its hydrophobicity, TPA^+^ elicits sizeable positive currents that reflect protein-independent interactions between the cationic substrate and the membrane (**Supplementary Figure 7**). However, for Gdx-Clo-7x, titration with mM concentrations of TPA^+^ also yields small negative capacitive currents that evolve more slowly than the TPA^+^ binding currents. These currents are consistent with electrogenic proton/TPA^+^ antiport. Peak current amplitudes, which reflect the initial rate of transport, were well fit by the Michaelis-Menten equation with a K_m_ value of 2 mM (**Figure 3B**). For comparison, EmrE transports TPA^+^ with a K_m_ value of 800 µM^18^. Gdm^+^ currents were nearly undetectable for the Gdx-Clo-7x mutant, but, as in EmrE, increasingly hydrophobic substitutions of the guanidinium moiety restored transport (**Figure 3C**). This trend differs from WT Gdx-Clo, for which hydrophobic substitution diminishes a substrate’s initial rate of transport^10^.

**Figure 3.**
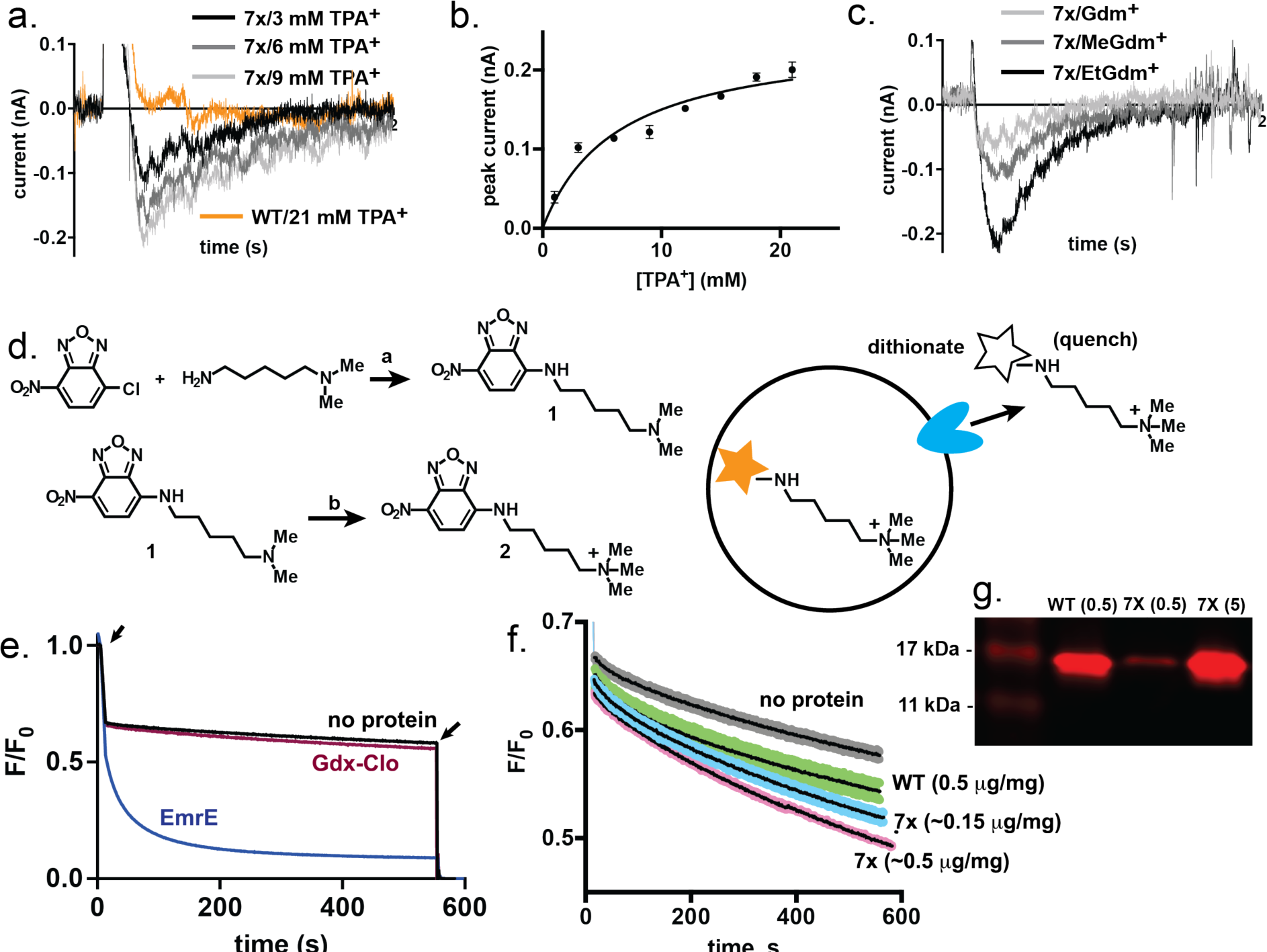
Gdx-Clo 7x transports quaternary ammonium compounds. A) Representative SSM electrophysiology traces for WT-Gdx-Clo (orange) or Gdx-Clo-7x (gray) upon perfusion with TPA^+^ at the concentrations indicated. Only perfusion with activating solution is shown. Control experiments with protein-free proteoliposomes are shown in **Supplementary Figure 7A**. B) Current amplitudes as a function of TPA^+^ concentration. Solid line represents a fit to the Michaelis-Menten equation with a K_m_ value of 2 mM. C) Representative SSM electrophysiology traces for Gdx-Clo-7x upon perfusion with 5 mM Gdm^+^, methylGdm^+^, or ethylGdm^+^. Only perfusion with activating solution is shown. Control experiments with protein-free proteoliposomes are shown in **Supplementary Figure 7B**. **D.** Left, two-step synthetic scheme for NBD-TA^+^. Reagents and conditions: (a) 4-chloro-7-nitrobenzo[c][1,2,5]oxadiazole (1 equiv), N^1^,N^1^-dimethylpentane-1,5-diamine (1 equiv), Et_3_N (2 equiv), DMF, 90 °C, 3 hrs, 48 %; (b) N^1^,N^1^-dimethyl-N5-(7-nitrobenzo[c][1,2,5]oxadiazol-4-yl)pentane-1,5-diamine (1 equiv), Et_3_N (0.1 equiv), Methyl iodide (5 equiv), DCM, rt, 1 hr, 100%. Right, cartoon of transport assay using NBD-TA^+^. Proteoliposomes with internalized NBD-TA^+^ (500 µM) are diluted into buffer containing 5 mM dithionite, which quenches external NBD-TA^+^. E) Representative timecourses of NBD-TA^+^ transport from liposomes containing no protein (black), WT Gdx-Clo (0.5 µg protein/mg lipid; maroon), or EmrE (0.5 µg protein/mg lipid; dark blue). Arrows show addition of dithionate (left) and Triton (right). Traces are normalized with respect to the fluorescence reading at 10 s. F) Timecourse of NBD-TA^+^ efflux from liposomes containing no protein, WT Gdx-Clo, or Gdx-Clo-7x. Dark line represents the mean of three independent replicates, and the standard error is shown by the colored regions (gray for no protein, green for WT-Gdx-Clo, and blue and pink for two concentrations of Gdx-Clo-7x.) The approximate protein density (µg protein/mg lipid), determined by quantitative Western blot, is shown in parentheses. Only the portion of the trace after dithionate addition and before Triton addition is shown. Full traces are shown in **Supplementary Figure 8**. G) Western blot analysis of reconstitution efficiency for WT Gdx-Clo (0.5 µg protein input/mg lipid) and Gdx-Clo-7x (0.5 or 5 µg protein input/mg lipid). Uncropped image is shown in **Supplementary Figure 9**. Quantification of band intensities from replicate measurements is shown in **Supplementary Table 2**.

We were unable to assess CTA^+^ transport using SSM electrophysiology, due to its detergent-like properties that cause it to partition into the membrane. To develop an assay for this antiseptic, we leveraged our structural observation that CTA^+^ binds with its alkyl tail extending out through a lateral portal in the protein and into the membrane, shown in Figure 2. This allowed us to adapt an approach used to study lipid transport by lipid transport proteins^30–32^, which bind substrate lipids in a similar manner^33^. We chemically synthesized a novel substrate analog with a fluorophore, nitrobenzoxadiazole (NBD), conjugated to an aliphatic 5-carbon linker with a trimethylammonium headgroup (NBD-TA^+^, **Figure 3D**). Based on our structure, if the quaternary ammonium headgroup binds similarly in the binding pocket, we expect the hydrophobic linker and NBD to extend through the portal and into the membrane, similar to the interaction between scramblases and NBD-conjugated lipids. For the assay, NBD-TA^+^ is incorporated into proteoliposomes. The addition of dithionate to the external buffer quenches the fluorescence of external NBD-TA^+^, including NBD-TA^+^ in the outer membrane leaflet, reducing the signal by about half. For proteoliposomes reconstituted with EmrE at a protein: lipid ratio such that each liposome contains approximately one transporter, fluorescence continues to decrease by 80% over ∼200 seconds, reflecting protein-mediated export of NBD-TA^+^. Addition of triton at the end of the experiment solubilizes the lipid vesicles, permitting complete quenching of the remaining protected NBD (**Figure 3E**). Control experiments showed that NBD by itself is not transported. In contrast, for protein-free liposomes, the fluorescence signal from internal NBD-TA^+^ remains steady after the initial quenching step. Likewise, as anticipated from the resistance assays, minimal NBD-TA^+^ transport is observed for WT Gdx-Clo proteoliposomes.

To assess NBD-TA^+^ transport by Gdx-Clo-7x, we performed this assay with protein at two protein:lipid ratios (**Figure 3F**). Independent quantitative Western blots of proteoliposomes show that Gdx-Clo-7x is not incorporated into proteoliposomes as efficiently as WT Gdx-Clo (p = 0.046; **Figure 3G**). By increasing the protein input for the reconstitution, we were able to prepare Gdx-Clo-7x liposomes that contained a comparable amount of protein relative to the WT Gdx-Clo proteoliposomes (p = 0.55). At both protein: lipid ratios, Gdx-Clo-7x exhibits slow, but significant transport of NBD-TA^+^, which increases with the amount of protein (**Figure 3F**). Relative to the experiment with EmrE, Gdx-Clo-7x transports NBD-TA^+^ at a slower rate, and a relatively large proportion of substrate remained unquenched over the timecourse of the experiment, which may reflect low fractional activity of the reconstituted protein. Nevertheless, these experiments establish transport of NBD-TA^+^ substrate across the membrane by Gdx-Clo-7x. Together, the TPA^+^ and NBD-TA^+^ transport assays link the resistance phenotype to active, proton-coupled transport of quaternary ammonium substrates.

### The majority of mutations are essential for quaternary ammonium resistance

To assess whether all seven mutations introduced in Gdx-Clo-7x are essential for quaternary ammonium transport and resistance activity, or whether a subset of these mutations would be sufficient for this activity, we developed a modified Gibson assembly method to construct a combinatorial library of all 128 possible variants of the seven mutations (**Figure 4A**). This library was transformed into Δ*emrE E. coli* cells (at a ratio that ensured each bacterial cell possessed only one plasmid), plated (with or without antiseptic selection), and the pooled plasmid DNA isolated from the resulting colonies was analyzed using next generation sequencing (NGS; **Supplementary Table 3**). We elected to plate the bacteria to isolate any colonies with appreciable growth, rather than perform a competitive growth in liquid culture, so that any potential variant with low fitness would not be outcompeted during the growth. For each selection, we harvested a sufficient number of colonies, and obtained a sufficient number of sequencing reads, to ensure library coverage (see *Methods*)^34^.

**Figure 4.**
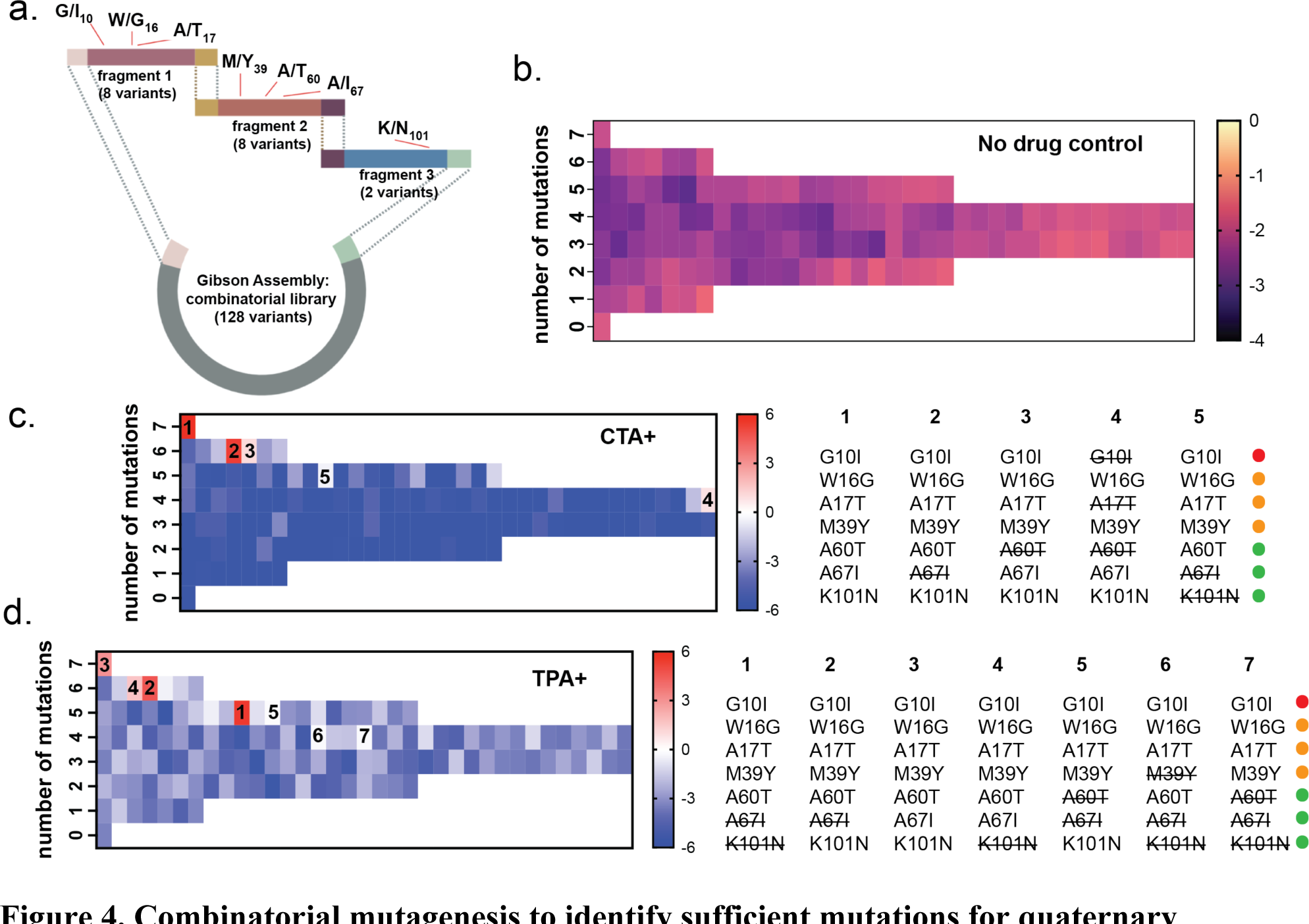
Combinatorial mutagenesis to identify sufficient mutations for quaternary ammonium transport. A) Schematic of the modified Gibson assembly method used to generate a combinatorial library of 128 mutational variants. B) Heatmap of library variants without selection (no drug control) assessed with NGS. Each box represents the reads for an individual variant as a fraction of total processed reads, displayed on a log_10_ scale. Values are the mean of three independent replicate experiments. Variants are grouped according to the number of mutations relative to WT Gdx-Clo (left axis). C, D) Enrichment coefficients (log_2_ scale) for library variants after selection with 120 µM CTA^+^ (C) or 18 mM TPA^+^ (D). The most highly represented variants are indicated (numbered in order of decreasing enrichment coefficient), with corresponding variants shown at right. Colored dots correspond to the mutational clusters from Figure 1. Values are derived from the mean of three independent replicate experiments (no-drug control and drug selection). Individual replicates are shown in **Supplementary Figure 10**. Correlations between replicate experiments are shown in **Supplementary Figure 11** and **Supplementary Table 4**. Enrichment coefficients for the CTA^+^ and TPA^+^ selections are shown in **Supplementary Tables 5** and **6**, respectively.

When this library was grown on non-selective media, all variants are well-represented with similar frequencies (**Figure 4B**, **Supplementary Figure 10**). This result shows that 1) all 128 variants are present in the library, and 2) that none of the variants are toxic to *E. coli* at the expression levels of this experiment. In contrast to the starting library shown in **Figure 4B**, selection with TPA^+^ or CTA^+^ eliminated or greatly reduced the frequency of most variants. Three independent replicates were performed for each growth condition, which showed good reproducibility with acceptable pairwise correlation coefficients above 0.71 for all comparisons, and >0.86 for most comparisons (**Supplementary Figure 10**, **Supplementary Figure 11**, **Supplementary Table 4**).

The variants that are enriched by the selections possess most of the originally identified seven mutations. Indeed, the most frequent variant isolated from the CTA^+^ screen possesses all seven, and variants lacking only A60T, A67I, or both were also enriched (**Figure 4C**). For TPA^+^ resistance (**Figure 4D**), variants lacking A67I, K101N, or both were enriched relative to the original library. Together, these experiments suggest that all the cluster 1 and 2 mutations (G10I, W16G, A17T, and M39Y; see Figure 1) are essential to confer robust quaternary ammonium resistance. While the cluster 3 mutations (A60T, A67I, and K101N) are, in general, less critical for this activity, they nonetheless contribute to fitness.

### Cluster 1 and 2 mutations introduce the major functional traits of SMR_Qacs_

The selection experiments above report on bacterial fitness, an amalgamate measure of quaternary ammonium transport along with other factors like protein folding and membrane insertion. To dissect the contribution of the three mutational clusters to substrate binding and export specifically, we purified proteins representing each of the mutant clusters (cluster 1: G10I; cluster 2: W16G, A17T, M39Y; cluster 3: A60T, A67I, K101N), as well as additional variants with combinations of mutations suggested by our NGS analysis. All of these mutants exhibited monodisperse size exclusion profiles, indicating proper folding **(Supplementary Figure 12)**. Tryptophan fluorescence quenching experiments showed that Gdm^+^ binding was impaired by both the cluster 1 (no detectable binding) and cluster 2 (K_d_ ∼10 mM) mutations. In contrast, the cluster 3 mutations resulted in a >10-fold *decrease* in Gdm^+^ K_d_ relative to WT Gdx-Clo (**Figure 5A**), perhaps due to the altered electrostatic environment conferred by K101N. For this cluster 3 mutant, we next evaluated the apparent K_d_ for Gdm^+^ over a range of pH values, in order to estimate the pK_a_ of the central glutamates. We found that the tighter Gdm^+^ binding of the cluster 3 mutant is accompanied by ∼5-fold weaker proton binding (pK_a_ of 7.1, compared to a pK_a_ of 6.7 for WT Gdx-Clo^9^; **Figure 5B**), suggesting a substantial perturbation to the Gdm^+^/H^+^ competition necessary for antiport. Indeed, SSM electrophysiology shows that electrogenic Gdm^+^/H^+^ antiport is eliminated in the cluster 3 mutant, as in the cluster 1 and 2 mutants that fail to bind substrate (**Figure 5C**). The cluster 2 transporter, however, retains the ability to transport ethyl- and phenylGdm^+^. This trend recapitulates the behavior of Gdx-Clo-7x and EmrE and contrasts with WT Gdx-Clo, which exhibits decreased transport currents as the hydrophobicity of the guanidinyl substrate is increased^10^.

**Figure 5:**
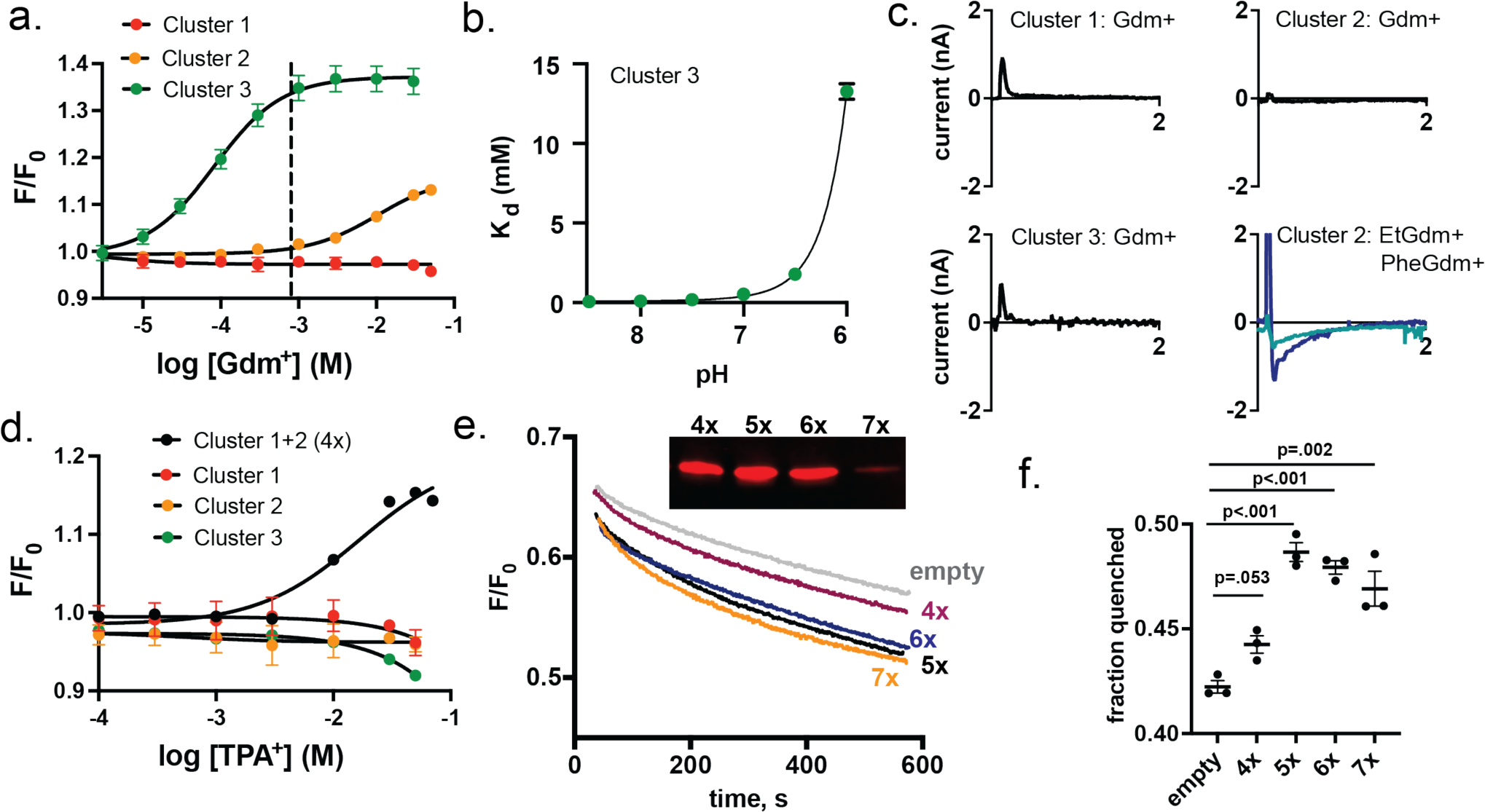
Functional characterization of the three variant clusters. A) Change in tryptophan fluorescence upon Gdm^+^ titration for cluster mutants as indicated. Datapoints represent the mean and SEM of three independent experiments. Solid lines represent fits to a single-site binding model with a K_d_ value of 80 μM for the cluster 3 mutant, and a K_d_ value of 10 mM for the cluster 2 mutant. The dashed line represents the K_d_ value for WT Gdx-Clo determined in Figure 2. B) Gdm^+^ K_d_ values measured as a function of pH for the cluster 3 mutant. The solid line represents a fit to Eq. 1, with a K_d_ of 88 μM and a composite pK_a_ of 7.1. Datapoints are the mean and SEM of three independent experiments. C) SSM electrophysiology traces for indicated cluster mutants upon perfusion with the indicated substrates (10 mM Gdm^+^ for the cluster 1 and cluster 2 mutants; 1 mM Gdm^+^ for the cluster 3 mutant; 5 mM for EtGdm^+^ and PheGdm^+^). Only perfusion with activating solution is shown. D) Change in tryptophan fluorescence upon TPA^+^ titration for cluster mutants as indicated. Datapoints represent the mean and SEM of three independent experiments. For the cluster 1+2 mutant, the solid line represents a fit to a single-site binding model with a K_d_ value of 21 mM. E) Representative traces of NBD-TA^+^ transport by mutants 4x, 5x, 6x, and 7x. For clarity, only the transport phase is shown (between dithionate addition and detergent addition). The inset shows a Western blot of liposomes reconstituted with protein used for the transport assay. The uncropped blot is shown in **Supplementary Figure 13**. F) Summary of three independent NBD-TA^+^ transport experiments for mutants indicated. Datapoints represent the fraction of NBD quenched at 560 s, prior to detergent addition, and error bars represent the SEM. The differences between 5x, 6x, and 7x are not statistically significant (5x vs 6x p=0.66; 5x vs. 7x p=0.16; 6x vs. 7x p=0.47).

When introduced individually, none of the clusters permitted TPA^+^ binding. However, cluster 1 and 2, introduced together, were sufficient for TPA^+^ binding with a K_d_ of 21 mM, similar to that of Gdx-Clo-7x (**Figure 5D**). However, we are unable to observe TPA^+^ transport by this mutant using SSM electrophysiology. We note that this variant was relatively highly represented in our NGS sequencing, but still ∼10-fold less prevalent than Gdx-Clo-7x (**Supplementary Table 4**). Since TPA^+^ transport by Gdx-Clo-7x is only just above the threshold for detection by SSM electrophysiology (**Figure 3**), we expect that this cluster 1 + cluster 2 mutant, and other surviving variants, might contribute to low levels of TPA^+^ transport yet not meet the lower limit for electrophysiological detection.

CTA^+^, with its small headgroup, does not require any additional mutations to support binding to WT Gdx-Clo. However, our NGS experiments show that this same complement of cluster 1 and cluster 2 mutations is essential for CTA^+^ resistance. Indeed, weak transport is detected for the construct with the cluster 1 and 2 mutations (labelled 4x in Figure 5E). The introduction of additional cluster 3 mutations (Gdx-Clo-5x: A60T, Gdx-Clo-6x: A60T and K101N, and Gdx-Clo-7x: A60T, A67I, K101N) further increases the rate of NBD-TA^+^ export (**Figure 5E**, **F**). Although the measured differences in NBD-TA^+^ efflux from proteoliposomes reconstituted with Gdx-Clo-7x, Gdx-Clo-6x, and Gdx-Clo-5x are not significant, Western blot analysis of proteoliposomes used for this assay showed that Gdx-Clo-7x is incorporated ∼3-fold less efficiently than the other Gdx-Clo variants (**Figure 5E**, **inset**), suggesting that, per protein, Gdx-Clo-7x has the highest activity. Combining the results of the NGS analysis and the biochemical analysis, we find that cluster 1 and cluster 2 mutations are sufficient to introduce polyselective quaternary ammonium transport to the selective Gdx-Clo WT background, but that the introduction of additional mutations from cluster 3 increases the rate of quaternary ammonium export, contributing to increased fitness of the bacteria.

## Discussion

The goal of this work was to identify a minimal set of mutations that convert a selective member of the SMR family, Gdx-Clo, into a promiscuous transporter of quaternary ammoniums. Because of the structural and sequence similarity between the SMR_Qac_ and SMR_Gdx_ subtypes, we reasoned that this would be a straightforward way to focus on the molecular features that are most critical for quaternary ammonium transport by drug exporters in this family, such as EmrE.

We identified three sensitive locations that contribute to transport of two structurally distinct quaternary ammonium compounds. The first is G10, which resides at the packing interface between helix 1 and helix 3. Based on the structures of Gdx-Clo and EmrE, we propose that the G10I mutation disrupts packing and contributes to the disorganized binding site hydrogen bond network observed in substrate-bound EmrE^18^. The second mutational cluster is W16G/A17T/M39Y, located in the immediate vicinity of the central glutamate. This group of mutations removes one hydrogen bond to the central glutamate (W16G) but provides two adjacent hydrogen bonding possibilities (A17T and M39Y). We propose that this also facilitates rotameric flexibility for the central glutamates. The third is A60T/A67I/K101N, which is located in a region of the protein that undergoes large conformational changes during the inward-to-outward-facing conformational transition. Based on their location in the structure, and the rates observed in the CTA^+^ transport assay, we propose that the cluster 3 mutations improve antiseptic resistance by accelerating the rate of conformational change. None of the positions identified in this work have been previously pinpointed as playing a role in polyselectivity or substrate preference in the SMRs. Notably, however, previous mutational screens of EmrE do show that differential changes in resistance to several drugs occur for mutations to all seven^21,22^.

Of these seven mutations, six were subject to our focus initially because they are broadly conserved in the SMR_Gdx_ or SMR_Qac_ subtype, but differ between subtypes. Thus, it seems likely that the mutations investigated here are representative of natural mutations that contributed to the evolutionary shift in substrate specificity. Although a seventh mutation identified through directed evolution, A60T, was required to achieve robust quaternary ammonium resistance on the mutant Gdx-Clo background used in our initial screens, this mutation is not essential for quaternary ammonium transport. In follow-up resistance assays, we observed that several antiseptic-resistant variants lacked A60T, along with other cluster 3 residues.

The essential residues for polyspecific quaternary ammonium transport are contributed by cluster 1 (G10I) and cluster 2 (W16G/A17T/M39Y) (**Figure 6A**). By itself, G10I is non-functional in all scenarios that we investigated, although it overexpresses and is monodisperse upon purification. G10I does not bind or transport any guanidinium or quaternary ammonium substrate. In contrast, the cluster 2 mutations are less severe when introduced on the WT Gdx-Clo background. Although this complement of mutations decreases the protein’s affinity for Gdm^+^ by >10-fold, the cluster 2 mutant remains capable of transporting substituted guanidinium ions with the same substrate hydrophobicity trend seen for EmrE^10^. More importantly, the cluster 2 mutations permit introduction of the otherwise-intolerable G10I.

**Figure 6.**
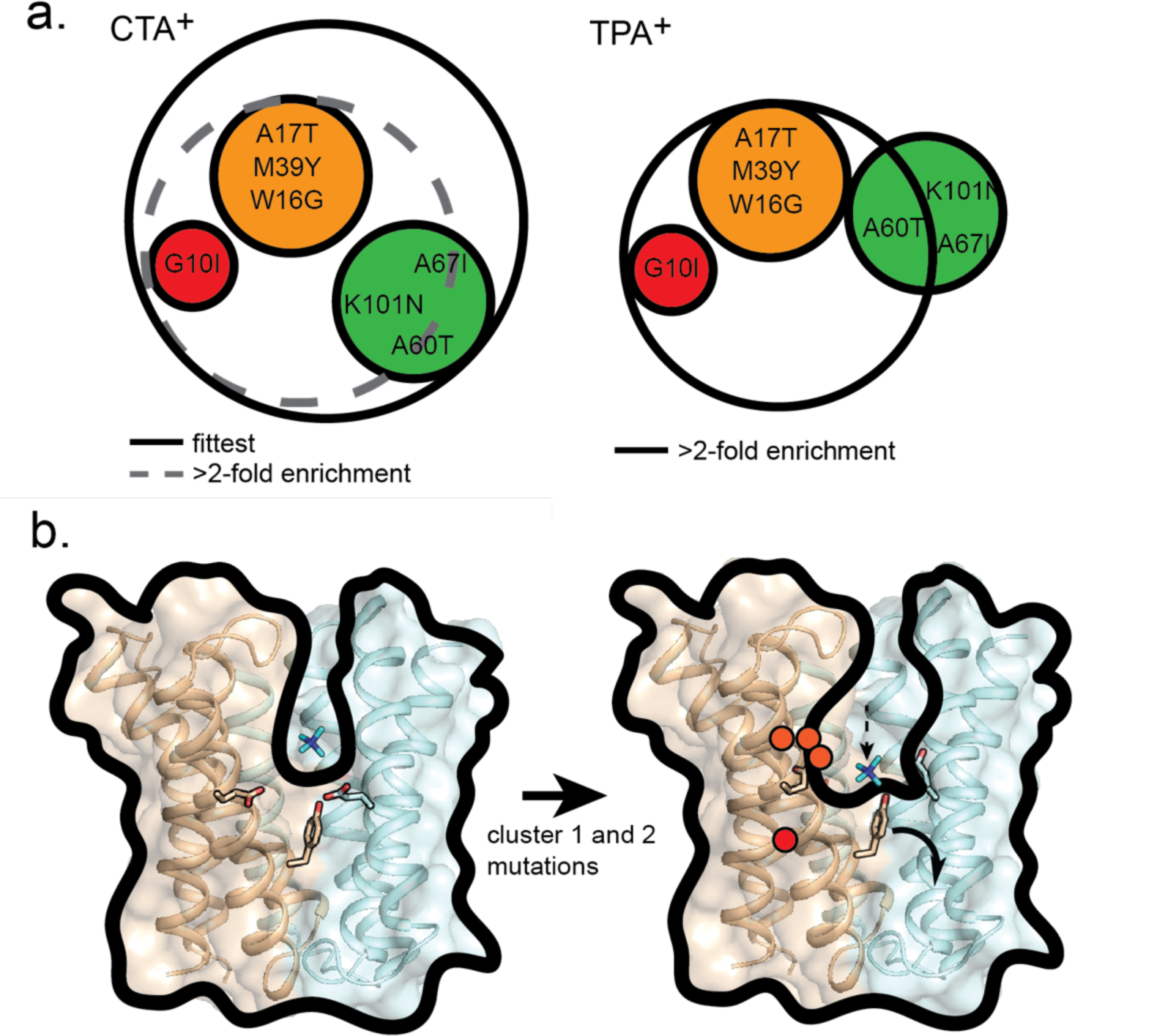
Summary of necessary and sufficient mutations to Gdx-Clo for quaternary ammonium resistance. A) For each antiseptic, mutations borne by the most highly enriched variant are indicated by the solid line. For CTA^+^, mutations borne by constructs that are enriched >2-fold are indicated by the dashed line. For TPA^+^, mutations encompassed by the solid line are present in all variants that are enriched >2-fold. B) Mechanistic model for the quaternary ammonium transport conferred by cluster 1 and 2 mutations. Left, in the WT transporter, a tight hydrogen bond network and rigid binding site prevent quaternary ammoniums from accessing the bottom of the pocket defined by the central glutamates (sticks). Right, the peripheral cluster 1 and 2 mutations allow more rotameric flexibility in binding site residues, which indirectly expands the binding pocket. Quaternary ammoniums penetrate between the central glutamates and trigger the tyrosine switch (sticks). Conformational change is accelerated by cluster 3 mutations (not shown).

The combination of G10I and the cluster 2 mutations meets our study’s minimal requirements for promiscuous quaternary ammonium transport (**Figure 6A**). Together, the four mutations enable binding of the bulky TPA^+^, TPA^+^ resistance, and CTA^+^ resistance and transport. We hypothesize that these mutations contribute to binding pocket flexibility, which not only permits TPA^+^ binding, but also allows CTA^+^ to penetrate more deeply into the aqueous vestibule. Whereas CTA^+^ hovers just above the central glutamates in the WT-like structure determined here, all transported substrates for both EmrE and Gdx-Clo reside in the plane of the central glutamates^9,10,18^. The substrates’ ability to disrupt binding between the glutamates a tyrosine “switch” has been proposed to trigger the conformational change essential for transport^18,35^. Thus, despite their differences in bulk, we propose that transport of TPA^+^ and CTA^+^ both originate in the loosening of the hydrogen bond network in the binding site, which allows sidechains to rearrange so that hydrophobic or bulky substrates can access the depths of the binding pocket (**Figure 6B**). In the Gdx-Clo-7x mutant, the gain of quaternary ammonium transport activity is accompanied by other traits characteristic of EmrE and other SMR_Qacs_, including the loss of Gdm^+^ transport activity and preference for additional substrate hydrophobicity^8^.

Our experiments also suggest a plausible evolutionary pathway between the selective SMR_Gdx_ subtype and the polyspecific SMR_Qac_s. In the evolution of quaternary ammonium transport by SMR proteins, it is unlikely G10I could have arisen early, as it abolishes transport function. And while the cluster 2 mutations do not eliminate transport of compounds like phenylGdm^+^, the protein loses its ability to transport the native metabolite Gdm^+^ without gaining a new physiologically relevant transport function. This implies that the early introduction of cluster 2 mutations is also not a viable evolutionary trajectory (although it is possible that an unknown natural substrate acted as a bridge). Indeed, protein evolution often requires accumulation of functionally neutral mutations that are permissive for later mutations that change the function^36^. It is possible that the cluster 3 mutations contributed to this – by increasing binding affinity to Gdm^+^, they may have offset the deleterious effect of cluster 2 mutations that, on their own, weaken Gdm^+^ binding. It is also possible that other permissive mutations arose during the evolution an ancestral SMR, and that these are not recapitulated by our mutation of extant SMR_Gdx_ Gdx-Clo.

In summary, we have established a set of necessary and sufficient residues that facilitate quaternary ammonium transport by SMR transporters. These residues are conserved among polyspecific SMR_Qacs_, and thus are likely to represent naturally important positions in model transporters like EmrE. Notably, these mutations are peripheral to the binding pocket, and the changes to specificity are not the result of mutations to residues that interact directly with the substrate. By zeroing in on a minimal set of mutations, we provide new insight into the essential molecular determinants of polyspecific transport, with implications for other promiscuous drug exporters.

## Methods

### Quaternary Ammonium Resistance Assays

Overnight cultures (12-16 hrs) of Δ*emrE E. coli* (Keio collection, Coli Genetic Stock Center, New Haven, CT) bearing the SMR genes in the pBAD24 vector were diluted to OD_600_ of 0.05 with LB media containing 13.3 mM arabinose to induce protein expression, 50 µg/mL kanamycin, and 100 µg/mL carbenicillin, then grown to OD_600_ 0.5 – 0.8 (37°C, 240 rpm). 10-fold serial dilutions were spotted onto plates containing 70 mM K_2_HPO_4_, pH 7.2, 0.2% arabinose, antibiotic, and 120 µM cetrimonium bromide or 18 mM tetrapropylammonium chloride, and examined for growth after 24-48 hours. Parallel control plates without quaternary ammonium compound were prepared for each experiment.

### Directed evolution

Directed evolution was performed on a background construct of Gdx-Clo with G10I, W16G, A17T, M39Y, A67I, K101N mutations in pBAD24 using the GeneMorph II EZClone Domain Mutagenesis Kit (Stratagene) with the manufacturer’s protocol and the following primers: <u>Forward Primer</u>: 5’-CAGGAGGAATTCACCATGGCGTGGCTGATC-3’ <u>Reverse Primer</u>: 5-ACAGCCAAGCTTATTAGCTGCTGGTCGCTTT-3’

The library was transformed into high-efficiency, electrocompetent DH5α cells and the transformants were collected to prepare a stock of the library plasmid DNA. For screening, the purified library was transformed into the Δ*emrE E. coli* strain via electroporation and used to inoculate an overnight growth. The overnight culture was diluted to OD_600_ of 0.05, and plasmid expression was induced with 0.2% arabinose. The culture was grown until the OD_600_ reached 0.5, and serial dilutions were plated on buffered LB with arabinose and, for selections, 200 µM-300 µM CTA^+^B, or 20 mM-30 mM TPA^+^. After 24 hours growth, plasmid DNA was isolated from individual colonies and sequenced.

### Combinatorial Library Construction, Selection Assays, and Illumina Sequencing

Gene blocks (**Supplementary Table 7**) were synthesized by Azenta (Chelmsford, MA) and prepared as a pooled master mix (10 ng/µL for each fragment). Fragments were assembled into the pBAD24 vector using HiFi DNA Assembly Kit (New England Biolabs, Ipswich, MA) with 1-hour, 50° C incubation and 0.2 pmol total fragments. After transformation, half of the recovery culture was plated on LB agar with carbenicillin to evaluate transformation efficiency, and the remaining half used to inoculate an overnight culture (LB with carbenicillin) for purification of the combinatorial plasmid library. The assembled combinatorial library was transformed into Δ*emrE E. coli* via electroporation, prepared for resistance assays as described above, and plated as 10-fold serial dilutions on LB with 120 µM cetrimonium bromide, 20 mM TPA^+^, or neither, without additional antibiotics. Plates with >1000 isolated colonies were resuspended, and the population plasmid DNA was isolated. This number of colonies corresponds to a >95% chance that there is a clone present for each of the 128 variants in the input library^34^. Illumina adapters were added via PCR using the following primers:

<u>Forward Illumina Adapter:</u> 5’-ACACTCTTTCCCTACACGACGCTCTTCCGATCTXXXX-3’ <u>Reverse Illumina Adapter:</u> 5’-GACTGGAGTTCAGACGTGTGCTCTTCCGATCTXXXX-3’ Next generation (Illumina) sequencing was performed by Azenta (Amplicon-EZ). For each experiment, >135,000 quality reads were obtained, corresponding to >1000-fold coverage of each variant in the unselected population, which is sufficient for a selection of this kind^37^. Processing included removal of reads <10. Three independent library preparations and selections were performed for each experiment. The enrichment coefficients for variant *i*, χ_i_, is given by:

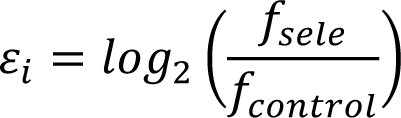

Where *f_sele_* and *f_control_* are given by the mean number of reads for the mutant (*N_l_*; mean of three experimental replicates), divided by the mean number of total reads (*N_tot_*) for the library, for libraries plated with and without quaternary ammonium compound, respectively.

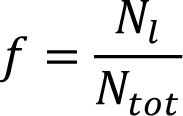

### Protein Purification and Liposome Reconstitution

Proteins were expressed and purified as previously described^8,10^. Briefly, hexahistidine-tagged proteins were overexpressed in *E. coli* (C43-DE3), extracted from *E. coli* membranes with 3% n-decyl-β-D-maltoside (DM; Anatrace) and purified using cobalt affinity resin and size exclusion chromatography. Purified protein was reconstituted into proteoliposomes (*E. coli* polar lipid extract, Avanti Polar Lipids, Alabaster, AL) by dialysis in 100 mM KCl, 100 mM K_2_HPO_4_, pH 7.52 as described^8,10^. For the NBD-TA^+^ transport assay, proteoliposomes were prepared with a protein:lipid ratio of 0.5 µg/mg. For SSM electrophysiology experiments, proteoliposomes were prepared with a protein: lipid ratio of 40 µg/mg. Proteoliposomes were stored at -80° C until use.

### Synthesis of NBD-TA^+^

All commercial reagents and solvents were used as supplied without further purification. Proton nuclear magnetic resonance (^1^H NMR) NMR spectroscopy was performed on a Bruker Advance 400 NMR spectrometer. ^1^H NMR shifts are reported in parts per million (ppm) and are referenced to tetramethylsilane (0.00 ppm) or residual DMSO (2.50 ppm). In the spectral data reported (**Supplementary Figure 14**), the format (δ) chemical shift (multiplicity, J values in Hz, integration) was used with the following abbreviations: s = singlet, d = doublet, t = triplet, q = quartet, p = pentet, m = multiplet. Electrospray ionization (ESI) mass spectral (MS) analysis was performed on a Thermo Scientific LCQ Fleet mass spectrometer. The final products were purified by reverse phase HPLC (RP-HPLC) with solvent A (0.1% of TFA in water) and solvent B (0.1% of TFA in CH_3_CN) as eluents with a flow rate of 45 mL/min. All final compounds have purity ≥95% as determined by Waters ACQUITY ultra-performance liquid chromatograph (UPLC) using reverse phase column (SunFire, C18, 5 μm, 4.6 × 150 mm2) and a gradient of solvent A (H_2_O with 0.1% of TFA) and solvent B (CH_3_CN with 0.1% of TFA) (**Supplementary Table 15**).

N^1^,N^1^-dimethylpentane-1,5-diamine (1): A solution of NBD-Cl (100mg, 0.5 mmol, 1 equiv.) in 0.5ml of anhydrous DMF was added dropwise in 5 min under a nitrogen atmosphere at room temperature, to a stirred solution of (5-aminopentyl) dimethylamine dihydrochloride (0.5 mmol, 1 equiv.) and triethylamine (154μl, 1 mmol, 2 equiv) in 1ml of anhydrous DMF. The reaction was stirred for 3 hours at 90 °C. The reaction was cooled to ambient temperature then subsequently purified by preparative reverse phase HPLC by direct injection of the reaction mixture using a gradient of 10%_50% B in 40 min_60ml. Product eluted at 26% B and the fractions were combined and lyophilized to produce compound **1** as the iminium (70mg, 0.25 mmol) as a mahogany-colored solid in 48% yield. ^1^H NMR (400 MHz, DMSO-*d*_6_) δ 10.30 (s, 1H), 9.55 (s, 1H), 8.51 (d, *J* = 9.0 Hz, 1H), 6.44 (d, *J* = 9.0 Hz, 1H), 3.49 (d, *J* = 7.6 Hz, 2H), 3.03 – 2.96 (m, 2H), 2.72 (s, 3H), 2.71 (s, 3H), 1.76 – 1.64 (m, 4H), 1.39 (p, *J* = 7.7 Hz, 2H).. ESI-MS m/z (M+H)^+^ = 294.16.

<u>N,N,N-trimethyl-5-((7-nitrobenzo[c][1,2,5]oxadiazol-4-yl)amino)pentan-1-aminium (2).</u> Triethylamine (2.3μl, 0.02 mmol) was added to a stirred solution of compound 1 (50mg, 0.17 mmol) in 3ml of DCM. Methyl iodide (53μl, 0.85 mmol, 5 equiv.) was then added and the reaction mixture turned a reddish-brown color. The reaction was stirred for 1 hour at ambient temperature. The reaction mixture was purified by preparative reverse phase HPLC by direct injection of the reaction mixture using the gradient of 10%_50% B in 40 min_60ml. Product eluted at 25% B and the fractions were combined and lyophilized to produce compound **1** (52mg, 0.17 mmol) as a mahogany-colored semi-solid in 100% yield. ^1^H NMR (400 MHz, DMSO-*d*_6_) δ 9.54 (s, 1H), 8.53 (d, *J* = 9.0 Hz, 1H), 6.44 (d, *J* = 9.0 Hz, 1H), 3.51 (overlapping, 2H), 3.28 – 3.23 (m, 2H), 3.03 (s, 9H), 1.78 – 1.68 (m, 4H), 1.38 (q, *J* = 7.7 Hz, 2H). ESI-MS m/z (M+H)^+^ = 308.24.

### NBD-TA^+^ transport

Proteoliposomes were preloaded with 500 µM NBD-TA^+^, subjected to 5 freeze/thaw cycles, and extruded 25 times through a 400-nm membrane filter. Liposomes were diluted 200-fold with assay buffer (100 mM KCl, 100 mM K_2_HPO_4,_ pH 7.5) containing 5 mM freshly prepared sodium hydrosulfite (dithionate) to quench external NBD-TA^+^. NBD-TA^+^ fluorescence was monitored (λ_ex_ = 470 nm, λ_em_ = 540 nm) using a fluorimeter (FP-8300, Jasco, Easton, MD) for 200 seconds prior to addition of 0.1% Triton X-100 to solubilize liposomes. Experiments were done in triplicate for each of two independent biochemical purifications.

*Tryptophan fluorescence binding assay:* Assay buffer was matched to the size exclusion chromatography buffer (100 mM NaCl, 10 mM HEPES, pH 8.0, 4 mM DM). Changes in tryptophan fluorescence upon substrate addition were monitored using a Jasco FP8300 fluorimeter (λ_ex_ = 280 nm, λ_em_ = 300-400 nm) and fit into a single-site binding isotherm. The pH dependence of the apparent K_d_ was fit to the following equation to determine the pK_a_ and K_d_:

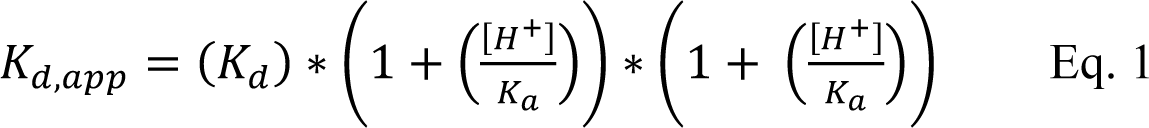

### Solid supported membrane (SSM) electrophysiology

SSM electrophysiology measurements were performed using the SURFE^2^R N1 (Nanion Technologies, Munich, Germany). Sensor preparation and verification of the sensor capacitance and conductance was performed exactly as described previously^10^. Experiments typically involved a series of three perfusions: a 2 s. perfusion with buffer, a 2 s. perfusion with activating solution (buffer plus substrate at the concentration indicated in the text), and a 4 s perfusion with buffer to wash away substrate. For a single sensor, currents from three perfusions with activating solution were averaged; these currents did not vary by more than 10%. Sensors were prepared from at least two independent protein purifications.

### Protein crystallization and x-ray crystallography

Purification of L10 monobodies and Gdx-Clo A60T were performed as previously described^10^. Freshly purified L10 and Gdx-Clo (10 mg/mL each) were mixed (2 monobody:1 transporter molar ratio) and combined with 50 µM CTA^+^ bromide prior to mixture with crystallization solution. Crystals formed at 22 °C in 100 mM CaCl_2_, 100 mM N-(2-acetamido)iminodiacetic acid (ADA) pH 6.75, and 33% PEG 600 and were frozen in liquid nitrogen prior to data collection at 13.5 keV at the Life Sciences Collaborative Access Team beamline 21-ID-D at the Advanced Photon Source, Argonne National Laboratory. Processed data was subjected to anisotropic truncation using Staraniso^38^, and phases were calculated by molecular replacement with Phaser^39^, with iterative rounds of refinement in Phenix^40^ and model building in Coot^41^. Polder omit maps were prepared using Phenix^42^.

## Acknowledgements

This work was supported by NSF CAREER award 1845012 to R.B.S. This research used resources of the Advanced Photon Source, a U.S. Department of Energy (DOE) Office of Science User Facility operated for the DOE Office of Science by Argonne National Laboratory under Contract No. DE-AC02-06CH11357. Use of the LS-CAT Sector 21 was supported by the Michigan Economic Development Corporation and the Michigan Technology Tri-Corridor (Grant 085P1000817).

## Data availability

Atomic coordinates for the crystal structures have been deposited in the Protein Data Bank under accession numbers 8VXU.

## Author Contributions

Conceptualization, O.E.B. and R.B.S.; Methodology, O.E.B., J.E.T, N.A.C., and R.B.S.; Investigation, O.E.B., E.O., J.H., E.M.G., I.R., R.M.L.; Writing – Original Draft, O.E.B. and R.B.S.; Writing – Review and Editing, O.E.B. and R.B.S.; Visualization, O.E.B., E.O., J.H., and R.B.S.; Supervision, R.B.S.; Project Administration, R.B.S.; Funding Acquisition, R.B.S.

## Competing interests

The authors declare no competing interests.

## Supplemental Material

Supplementary Figure 1. No-drug control for bacterial dilution assays in Figure 1A.

Supplementary Figure 2. Dilution assays for Gdx-Clo single mutant variants.

Supplementary Figure 3. FPLC profile for Gdx-Clo WT and Gdx-Clo-7x purification

Supplementary Figure 4. Representative SSM electrophysiology trace of Gdx-Clo A60T upon perfusion with 1 mM Gdm^+^

Supplementary Figure 5. Raw tryptophan fluorescence spectra for binding data shown in Figure 2.

Supplementary Figure 6. Second independent protein preparation of Gdx-Clo-7x: binding to Gdm^+^, CTA^+^, or TPA^+^, as indicated.

Supplementary Figure 7. Solid supported membrane electrophysiology: no protein controls.

Supplementary Figure 8. Full timecourse for the NBD-TA^+^ assays shown in Figure 3F.

Supplementary Figure 9. Uncropped Western blot for panel shown in Figure 3G.

Supplementary Figure 10. Heat maps showing raw counts for combinatorial library with no drug, CTA^+^ selection, and TPA^+^ selection.

Supplementary Figure 11. Correlations of enrichment coefficients (log_2_) between replicate selection experiments.

Supplementary Figure 12. FPLC profiles of mutant Gdx-Clo proteins used for functional analysis: cluster 1, cluster 2, cluster 3, cluster 1+2 (4x), 5x, 6x

Supplementary Figure 13. Uncropped Western blot from Figure 5E.

Supplementary Figure 14. Liquid chromatography/mass spectrometry analysis of NBD-TA^+^ synthesis.

Supplementary Figure 15. NMR analysis of NBD-TA^+^ synthesis

Supplementary Table 1. Data collection and refinement statistics for Gdx-Clo bound to CTA^+^.

Supplementary Table 2. Quantification of reconstitution efficiency measured by Western blot.

Supplementary Table 3. Sequencing statistics for replicate NGS experiments.

Supplementary Table 4. Correlation between enrichment coefficients of replicate selection experiments.

Supplementary Table 5. Most highly represented variants in CTA^+^ selection.

Supplementary Table 6. Most frequent variants in TPA^+^ selection. Fold change is shown relative to no-drug control experiments.

Supplementary Table 7. Gene block sequences for combinatorial library assembly.

---

**Supplementary Figure 1.**
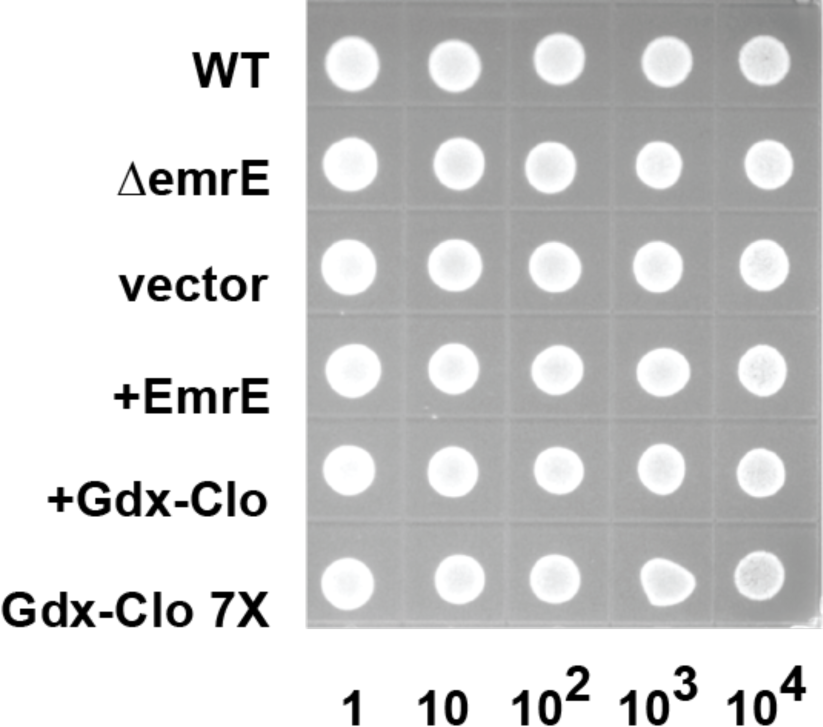
No-drug control for bacterial dilution assays in Figure 1A. Dilutions are indicated on the x-axis.

**Supplementary Figure 2.**
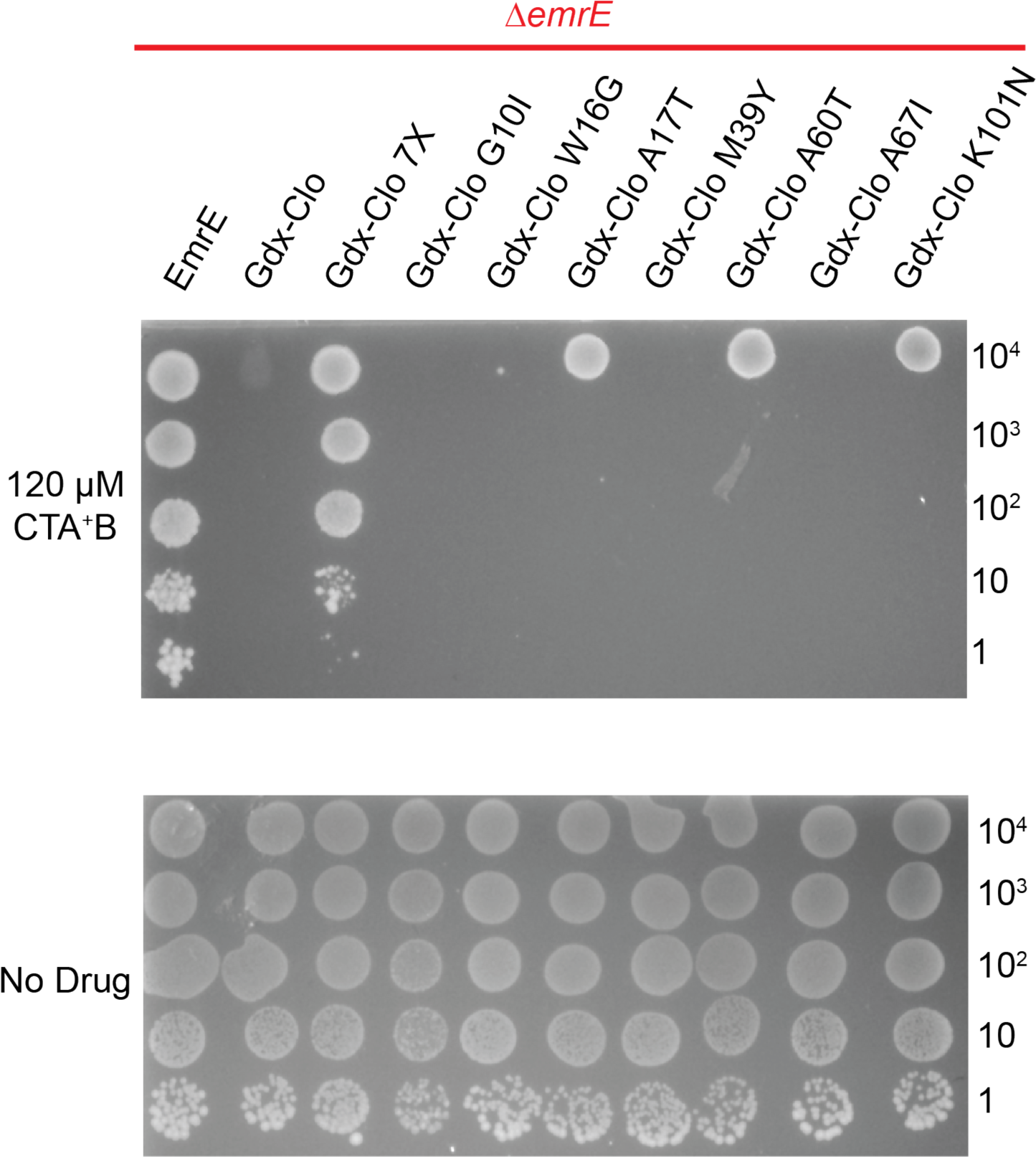
Dilution assays for Gdx-Clo single mutant variants. Dilutions are indicated on the y-axis.

**Supplementary Figure 3.**
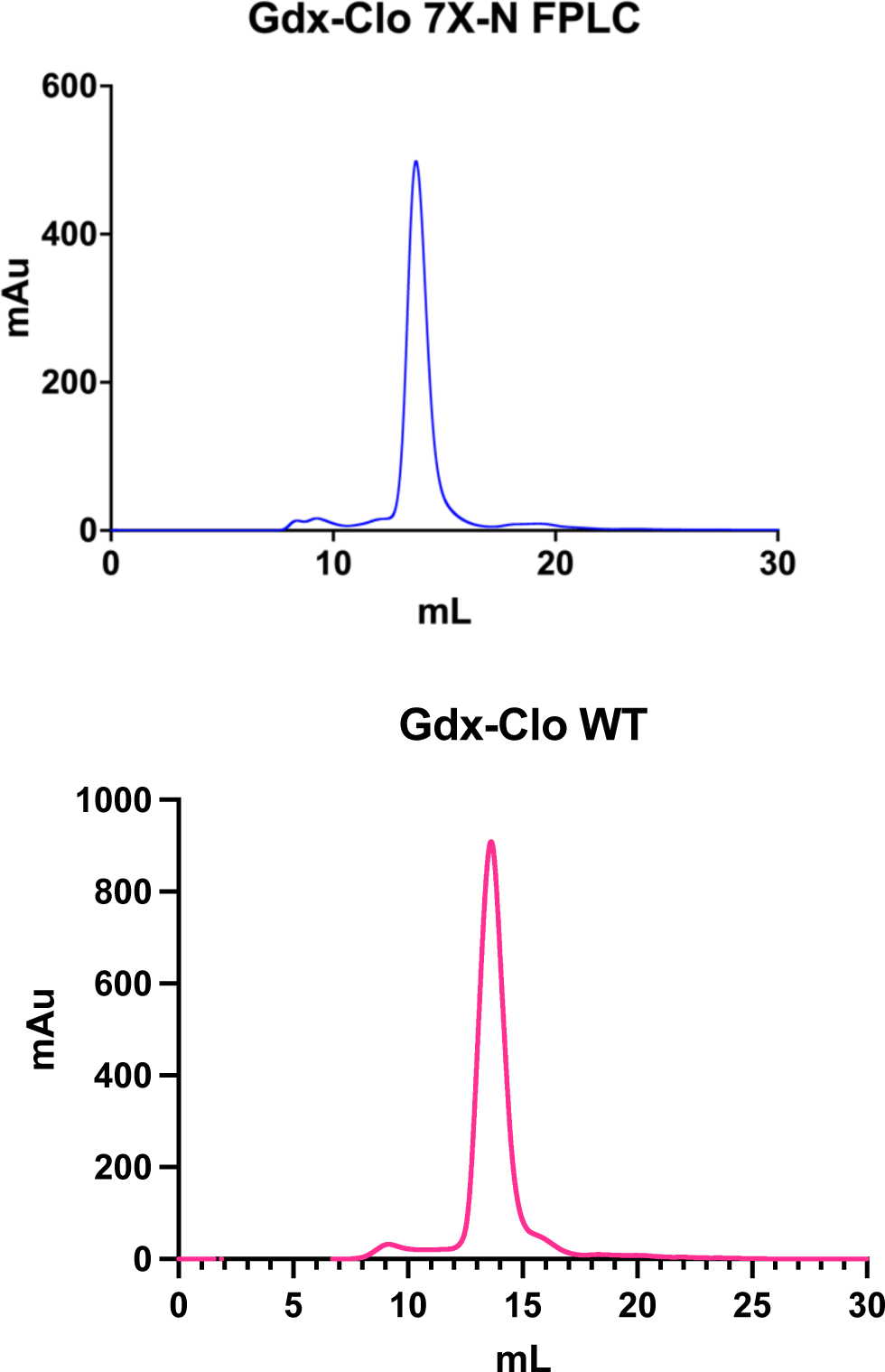
FPLC profile for Gdx-Clo WT and Gdx-Clo-7x purifications.

**Supplementary Figure 4.**
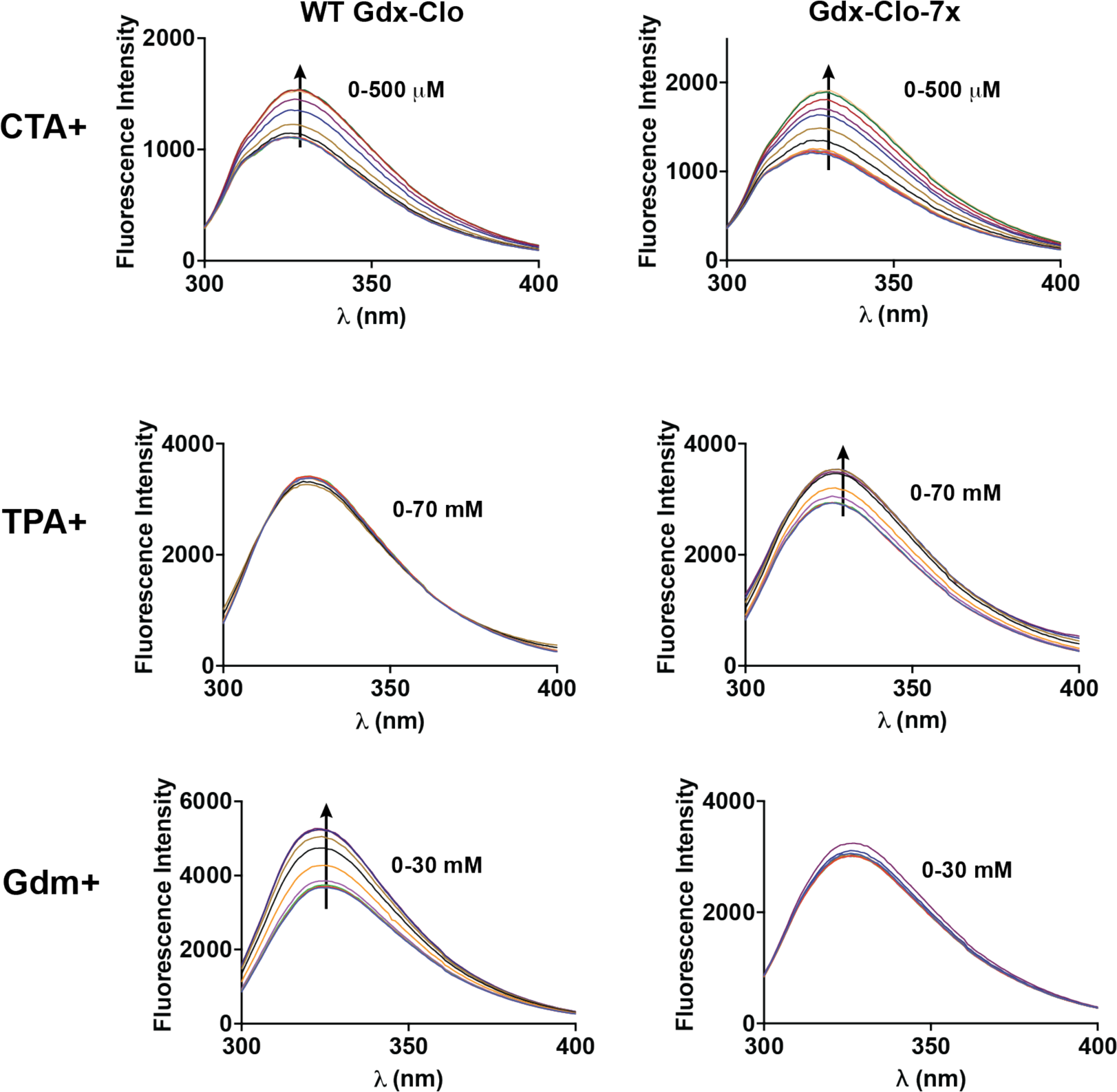
Raw tryptophan fluorescence spectra for binding data shown in Figure 2. Arrows indicate the direction of the change in fluorescence with increasing substrate.

**Supplementary Figure 5.**
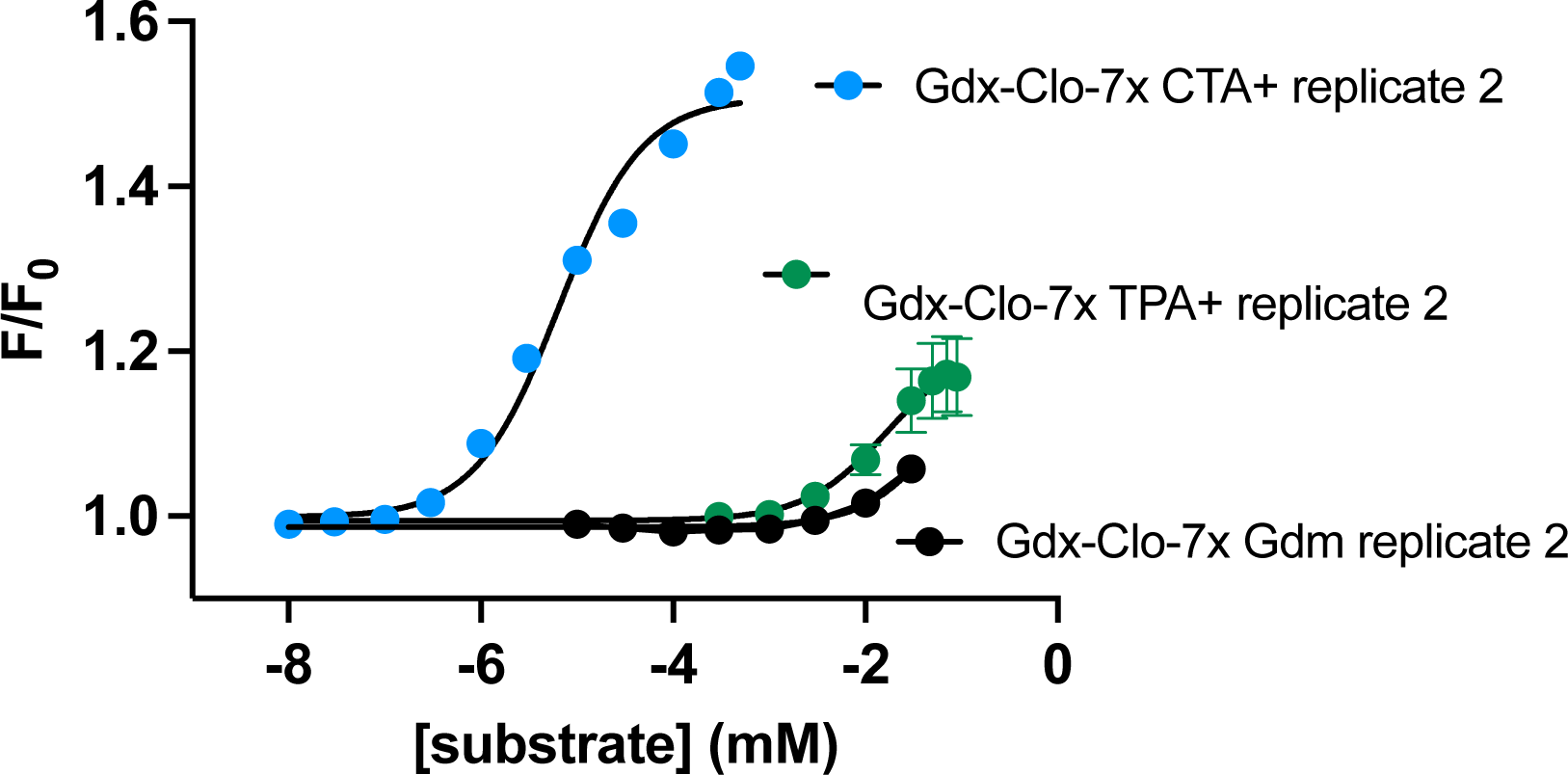
Second independent protein preparation of Gdx-Clo-7x: binding to Gdm^+^, CTA^+^, or TPA^+^, as indicated. Solid lines show fits to a single site binding isotherm with K_d_ values of 6.5 μM for CTA^+^ and 17.5 mM for TPA^+^.

**Supplementary Figure 6.**
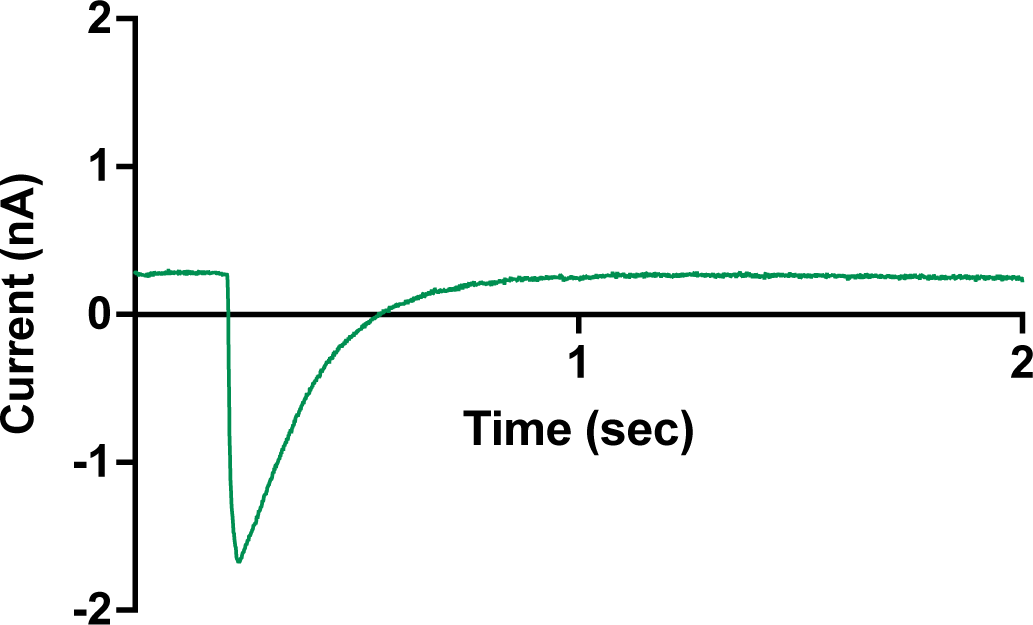
Representative SSM electrophysiology trace of Gdx-Clo A60T upon perfusion with 1 mM Gdm^+^. Only the perfusion with activating solution is shown. Trace is representative of experiments from two independently prepared sensors.

**Supplementary Figure 7.**
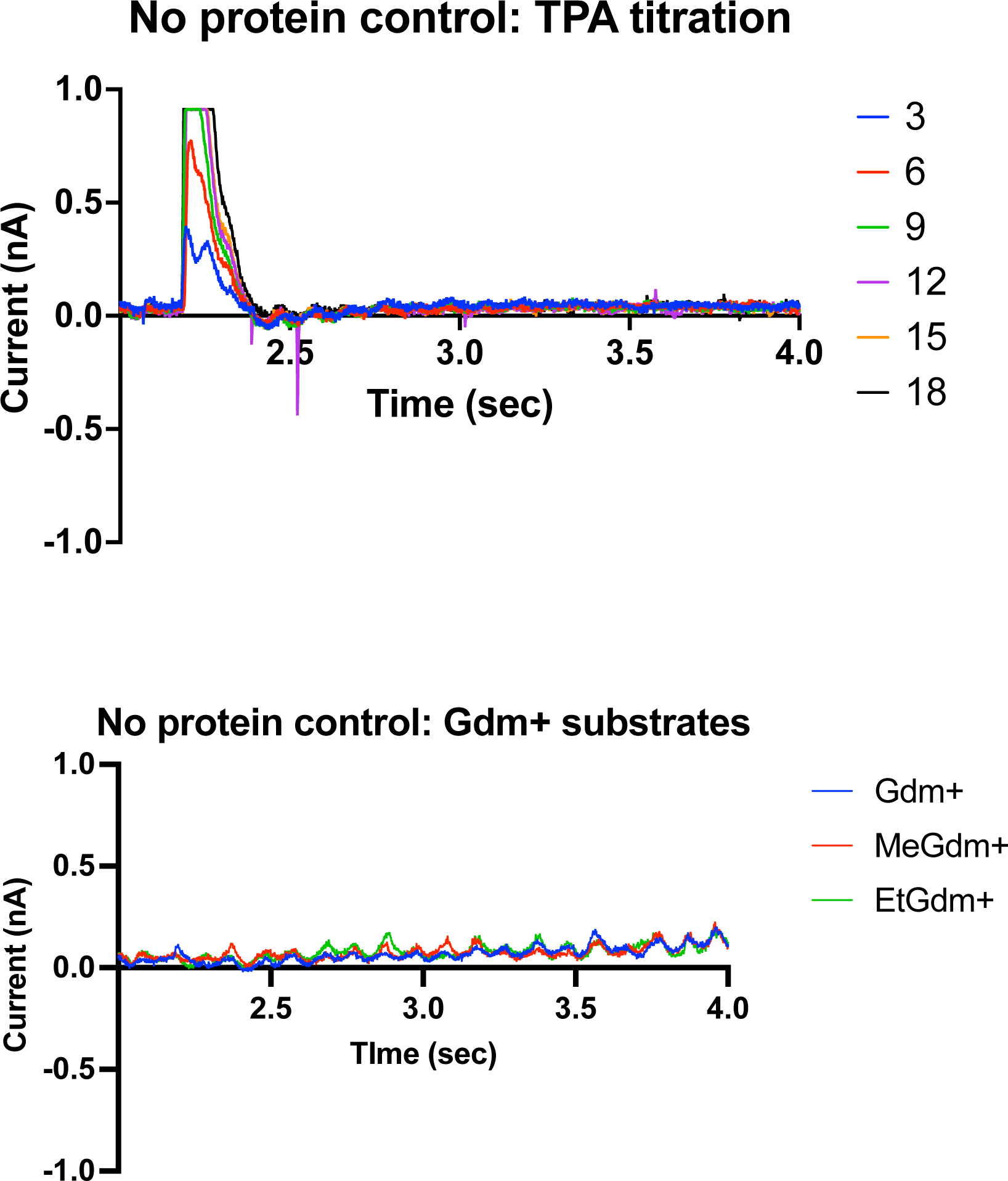
Solid supported membrane electrophysiology: no protein controls. TPA^+^ was titrated from 3-18 mM as indicated. Gdm^+^ and substituted guanidiniums were perfused at 1 mM.

**Supplementary Figure 8.**
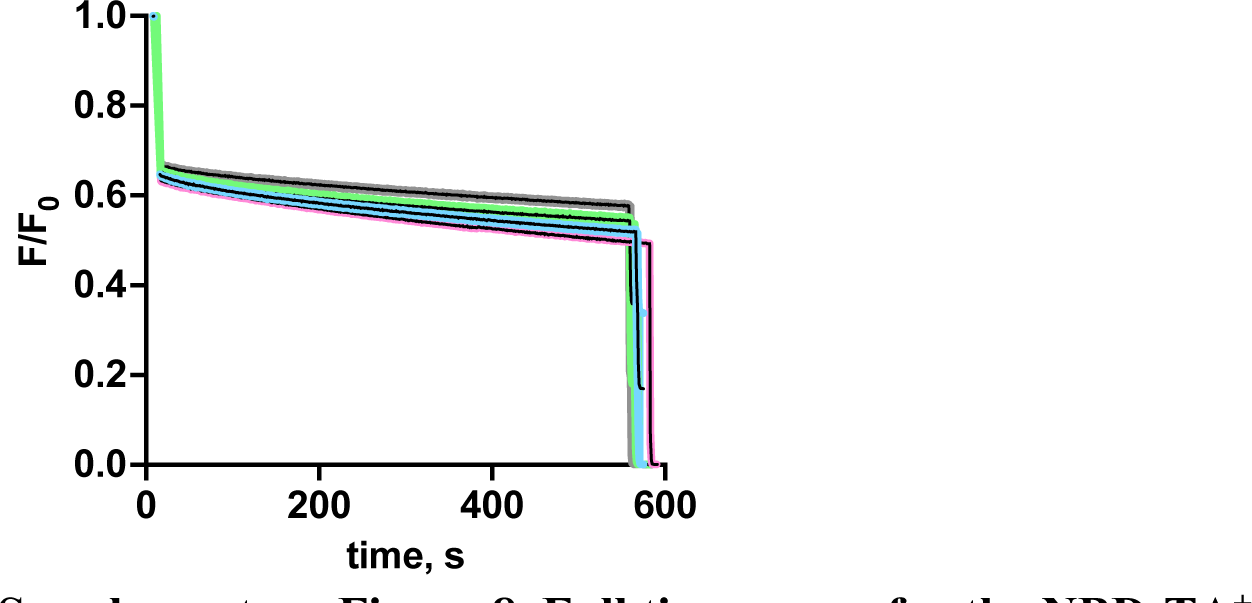
Full timecourse for the NBD-TA^+^ assays shown in Figure 3f. Black line shows the mean of three independent replicates. The error of the replicate measurements is shown by the colored regions (gray for no protein, green for WT Gdx-Clo, blue for Gdx-Clo-7x at low protein concentration, and pink for Gdx-Clo-7x at high concentration).

**Supplementary Figure 9.**
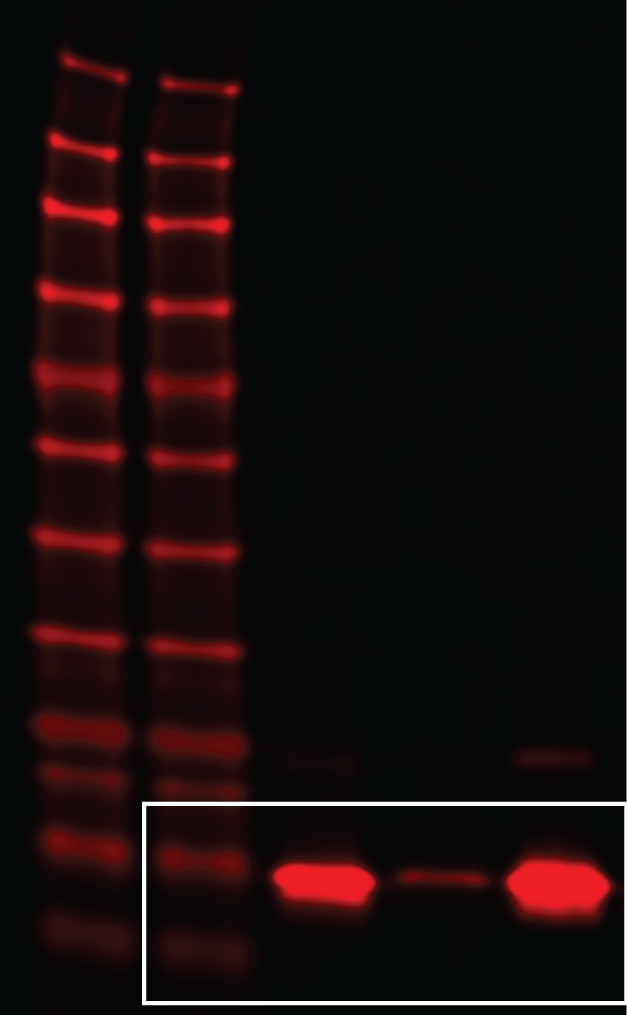
Uncropped Western blot for panel shown in Figure 3G. The portion used for Figure 3G is shown by the white boxed area.

**Supplementary Figure 10.**
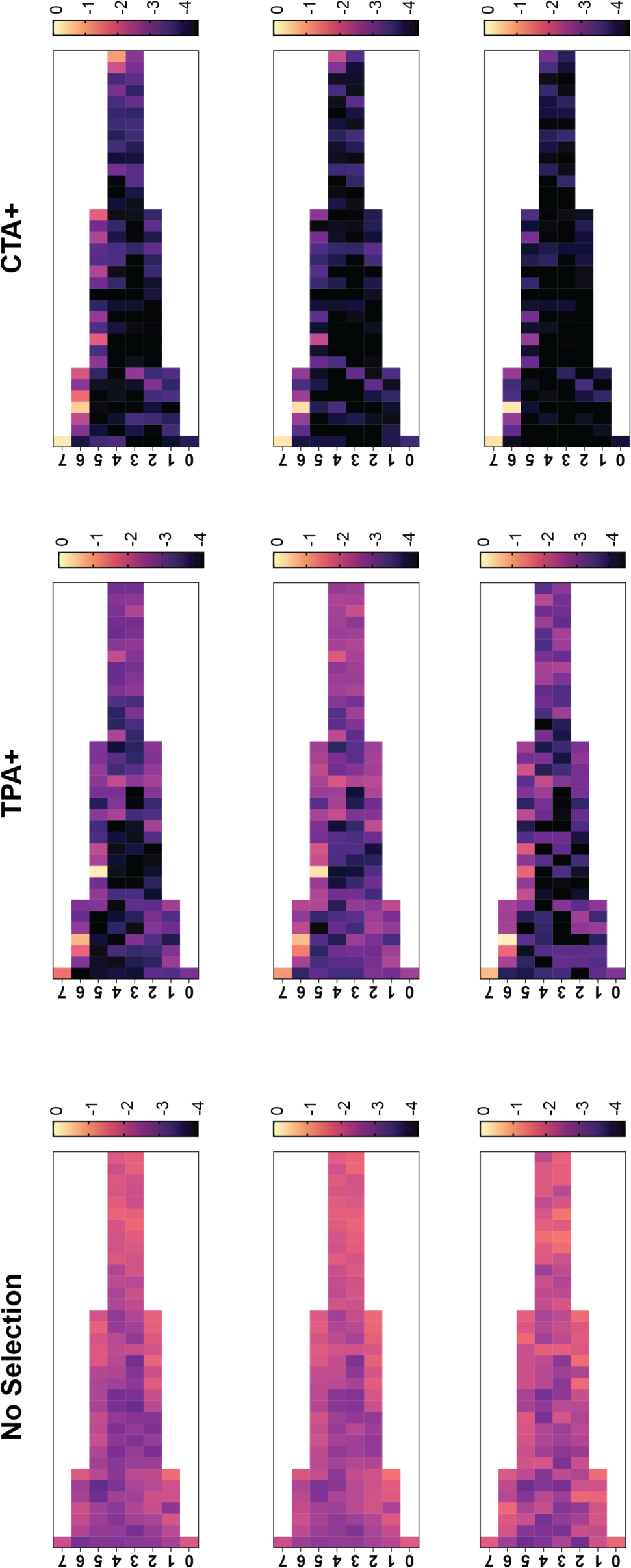
Heat maps showing raw counts for combinatorial library with no drug (also shown in Figure 4), CTA^+^ selection, and TPA^+^ selection. Each box represents reads for an individual variant divided by total reads. Data are visualized on a log_10_ scale.

**Supplementary Figure 11.**
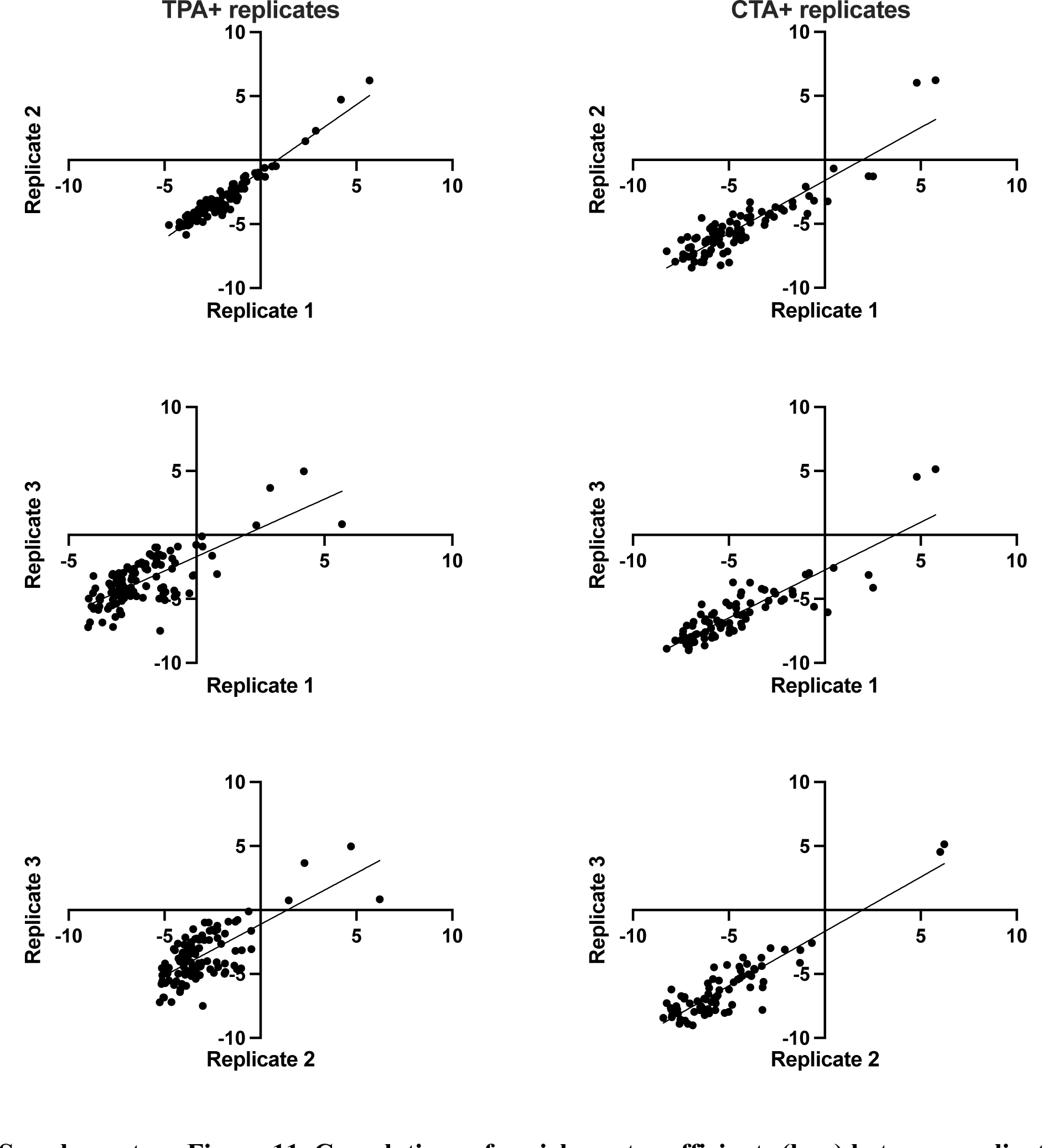
Correlations of enrichment coefficients (log_2_) between replicate selection experiments. Linear fits to the data are shown. R-values for fits are shown in Supplementary Table 4.

**Supplementary Figure 12.**
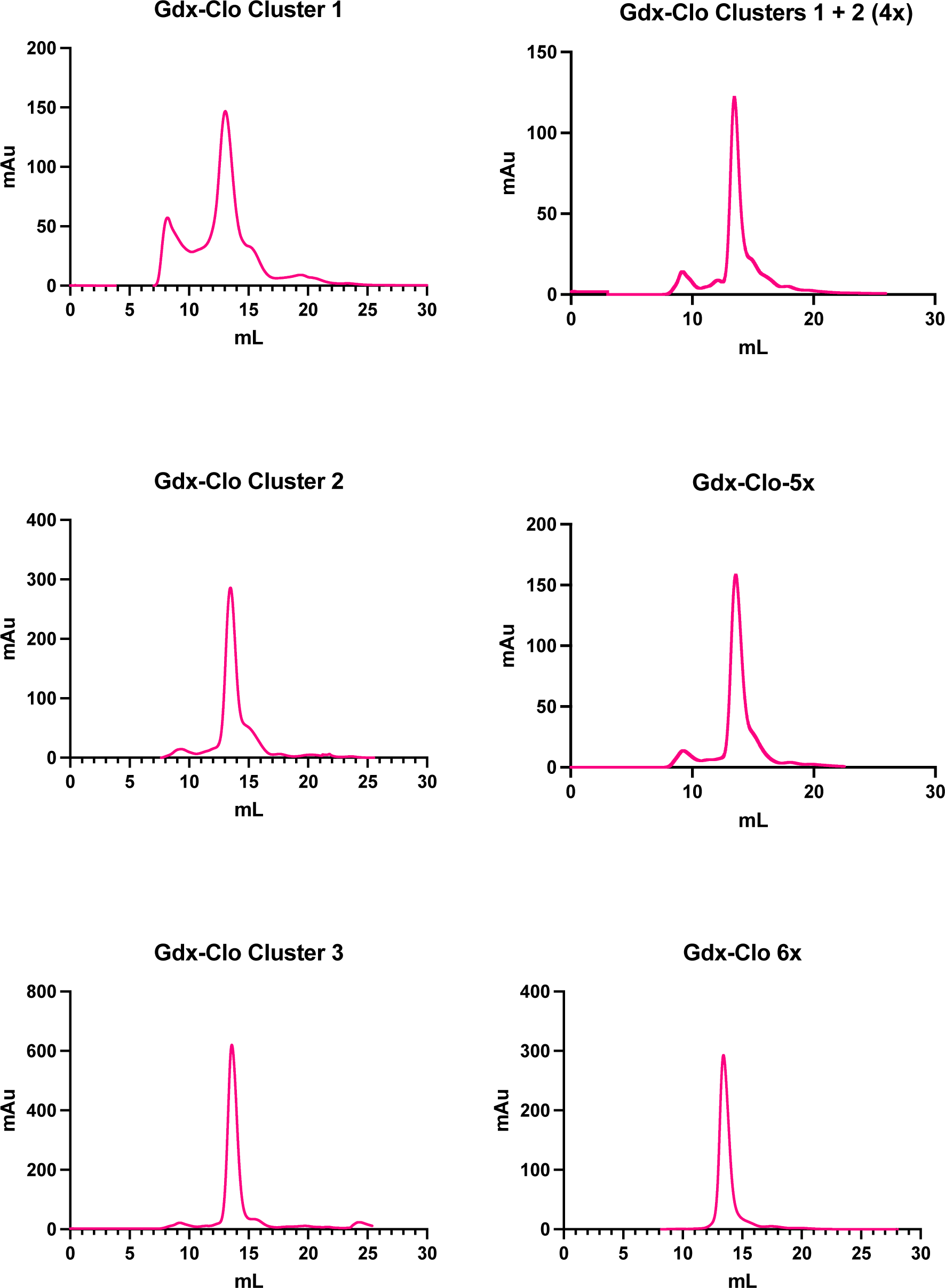
FPLC profiles of mutant Gdx-Clo proteins used for functional analysis: cluster 1, cluster 2, cluster 3, cluster 1+2 (4x), 5x, 6x.

**Supplementary Figure 13.**
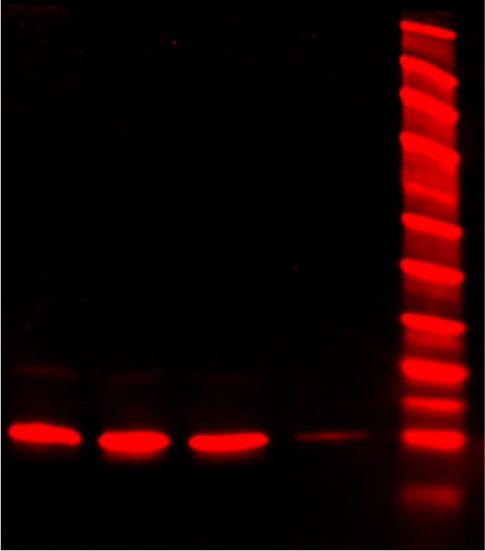
Uncropped Western blot from Figure 5E.

**Supplementary Figure 14.**
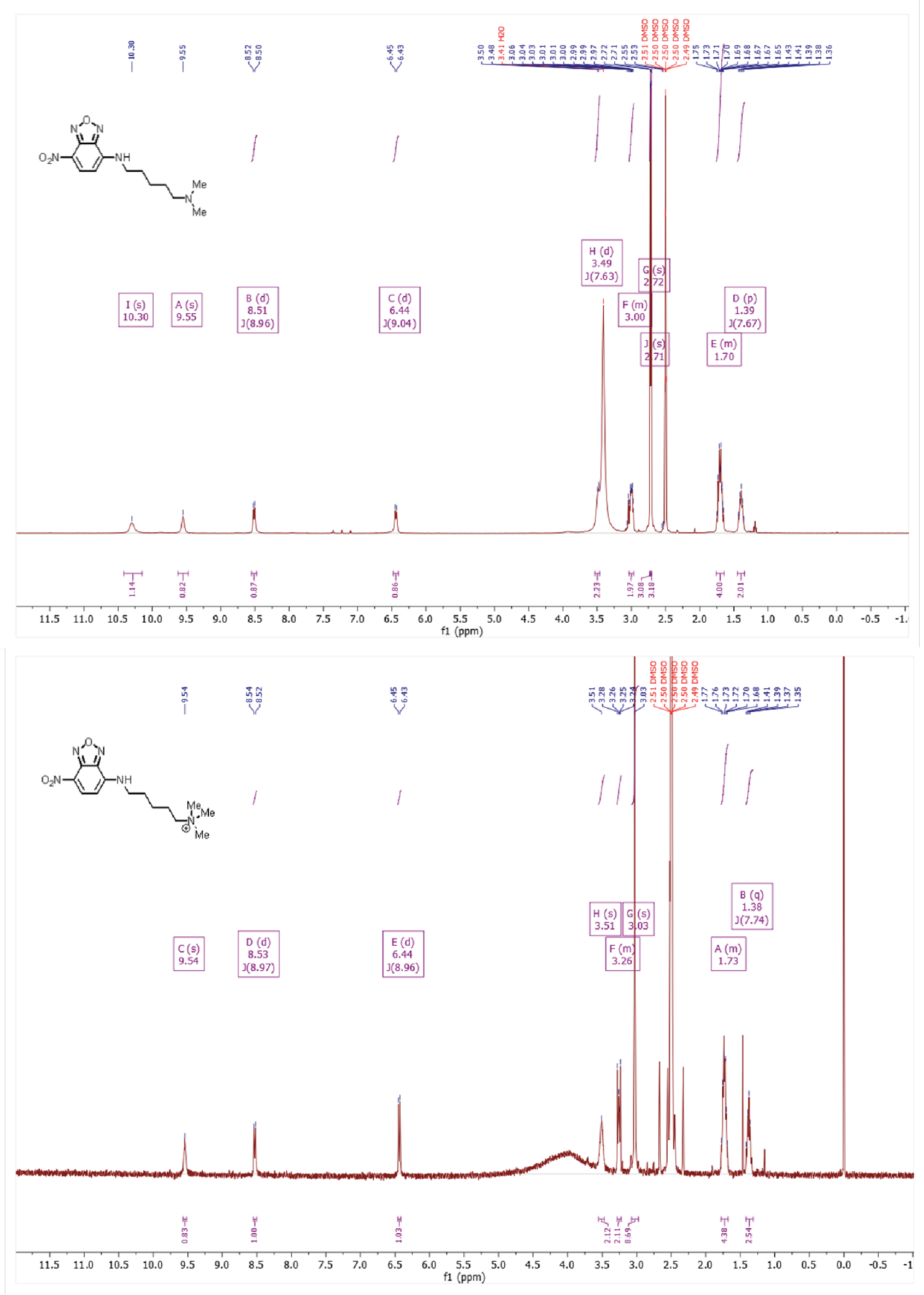
NMR analysis of NBD-TA^+^ synthesis

**Supplementary Figure 15.**
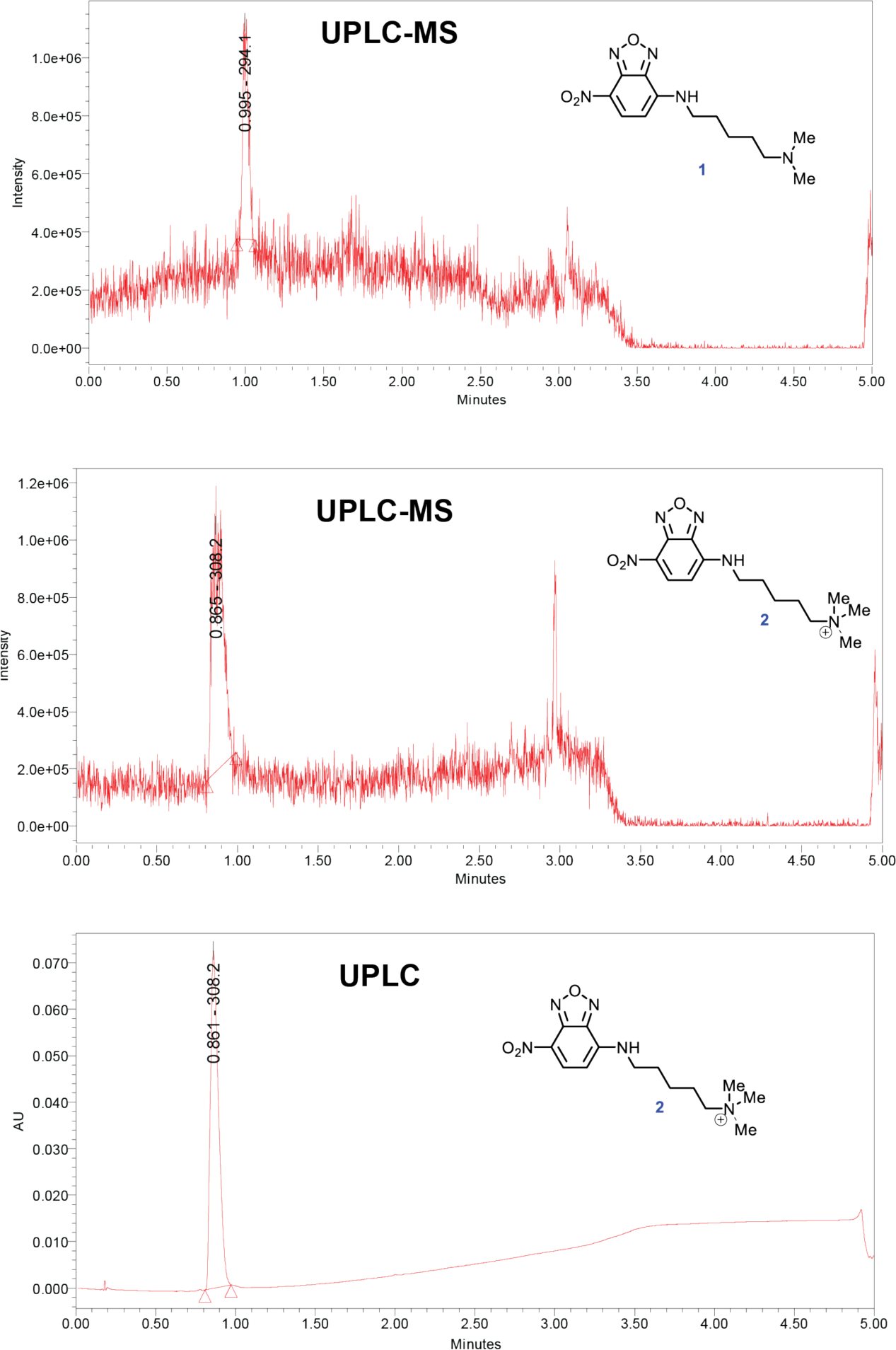
Liquid chromatography and mass spectrometry analysis of NBD-TAB synthesis.

**Supplementary Table 1.** Data collection and refinement statistics for Gdx-Clo A60T bound to CTA^+^.

|  |  |
| --- | --- |
| **Data Collection** |  |
| Space Group | P1 |
| Cell dimensions<br>*a*, *b*, *c*, (Å) | 50.13, 74.87, 108.25 |
| $\alpha$, $\beta$, $\gamma$ (°) | 93.27, 89.93, 109.51 |
| Resolution (Å) | 70.5-2.29 |
| Ellipsoidal Resolution Limit (best/worst) | 2.29/2.75 |
| % Spherical Data Completeness | 31.3 (4.0) |
| % Ellipsoidal Data Completeness | 82.1 (68.9) |
| *CC*<sub>1/2</sub> | 0.98 (.614) |
| Mn *I*/σ*I* | 6.2 (2.1) |
| Multiplicity | 3.8 (3.8) |
| **Refinement** |  |
| Resolution (Å) | 35.22 – 2.29 |
| No. reflections | 22,068 |
| R<sub>work</sub> / R<sub>free</sub> | 27.7/28.9 |
| Ramachandran Favored | 94.9 |
| Ramachandran Outliers | 1.1 |
| Clashscore | 10.6 |
| **R.m.s. deviations** |  |
| Bond lengths (Å) | 0.003 |
| Bond angles (°) | 0.667 |
| Coordinates in Protein Databank | 8VXU |

**Supplementary Table 2.** Quantification of Western blot band intensity. The difference between WT-Gdx-Clo (0.5) and Gdx-Clo-7x (5) is not significant (p = 0.41; t-test).

| | Protein input ($\mu\text{g}/\text{mg}$ lipid) | Mean ± SEM (AU) | n |
| --- | --- | --- | --- |
| WT Gdx-Clo | 0.5 | $23.5 \pm 2.9$ | 8 |
| Gdx-Clo-7x | 0.5 | $8.6 \pm 0.5$ | 5 |
| Gdx-Clo-7x | 5 | $27 \pm 2.9$ | 7 |

**Supplementary Table 3.**
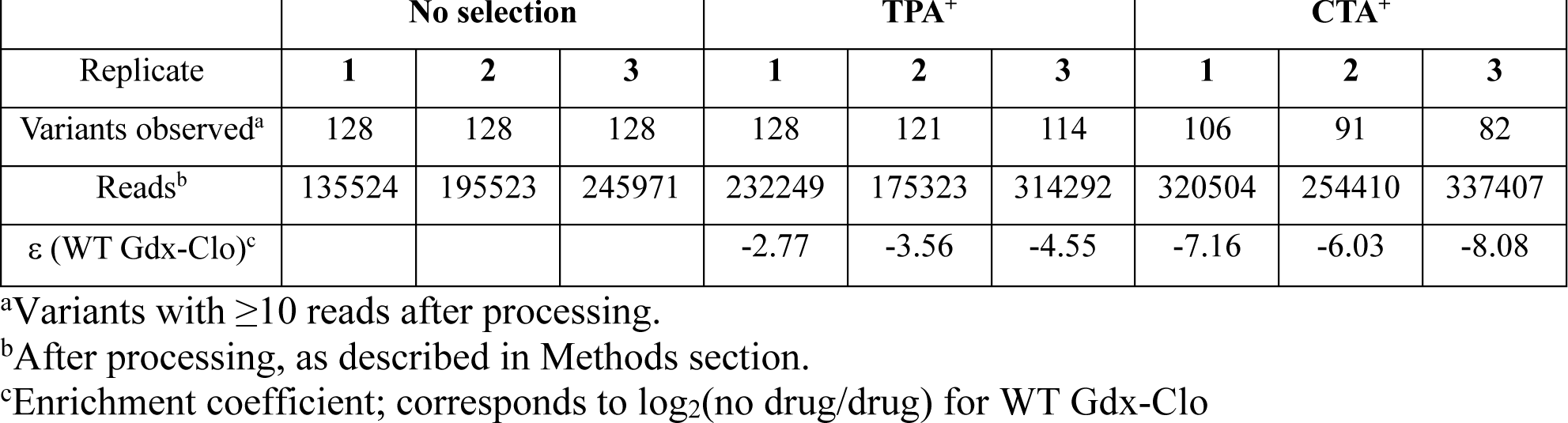
Sequencing statistics for replicate NGS experiments.

|  | No selection |  |  | TPA<sup>+</sup> |  |  | CTA<sup>+</sup> |  |  |
| --- | --- | --- | --- | --- | --- | --- | --- | --- | --- |
| Replicate | 1 | 2 | 3 | 1 | 2 | 3 | 1 | 2 | 3 |
| Variants observed<sup>a</sup> | 128 | 128 | 128 | 128 | 121 | 114 | 106 | 91 | 82 |
| Reads<sup>b</sup> | 135524 | 195523 | 245971 | 232249 | 175323 | 314292 | 320504 | 254410 | 337407 |
| $\varepsilon$ (WT Gdx-Clo)<sup>c</sup> | | | | -2.77 | -3.56 | -4.55 | -7.16 | -6.03 | -8.08 |

<sup>a</sup>Variants with $\geq 10$ reads after processing.
<sup>b</sup>After processing, as described in Methods section.
<sup>c</sup>Enrichment coefficient; corresponds to $\log_2(\text{no drug/drug})$ for WT Gdx-Clo

**Supplementary Table 4.**
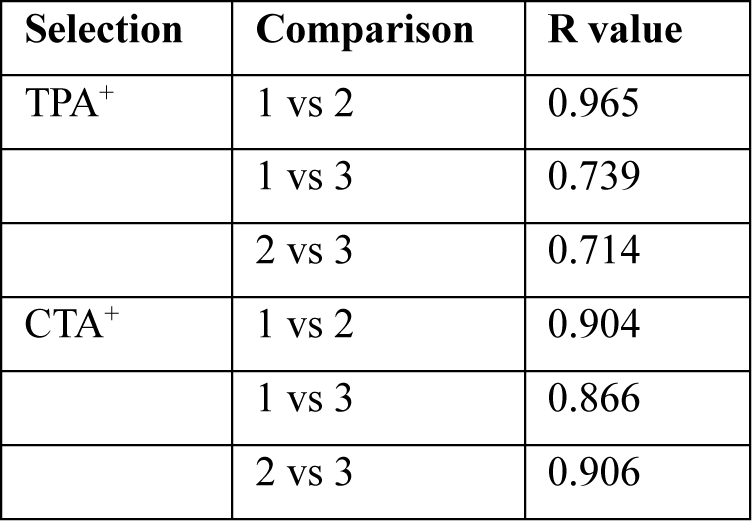
Correlation between enrichment coefficients of replicate selection experiments.

| Selection | Comparison | R value |
| --- | --- | --- |
| TPA<sup>+</sup> | 1 vs 2 | 0.965 |
|  | 1 vs 3 | 0.739 |
|  | 2 vs 3 | 0.714 |
| CTA<sup>+</sup> | 1 vs 2 | 0.904 |
|  | 1 vs 3 | 0.866 |
|  | 2 vs 3 | 0.906 |

**Supplementary Table 5.** Most highly represented variants in CTA^+^ selection. Change in frequency is shown relative to no-drug control experiments. All variants present at a frequency of at least 50% of the no-drug plates are shown. Colors correspond to mutational clusters described in Figure 1.

| Rank | Fold change | G10I | W16G | A17T | M39Y | A60T | A67I | K101N |
| --- | --- | --- | --- | --- | --- | --- | --- | --- |
| 1 | 55 | ● | ● | ● | ● | ● | ● | ● |
| 2 | 38 | ● | ● | ● | ● | ● |  | ● |
| 3 | 2 | ● | ● | ● | ● |  | ● | ● |
| 4 | 1.8 |  | ● |  | ● |  | ● | ● |
| 5 | 0.7 | ● | ● | ● | ● | ● |  |  |

**Supplementary Table 6.** Most frequent variants in TPA^+^ selection. Change in frequency is shown relative to no-drug control experiments. All variants present at a frequency of at least 50% of the no-drug plates are shown. Colors correspond to mutational clusters described in Figure 1.

| Rank | Fold change | G10I | W16G | A17T | M39Y | A60T | A67I | K101N |
| --- | --- | --- | --- | --- | --- | --- | --- | --- |
| 1 | 43 | ● | ● | ● | ● | ● |  |  |
| 2 | 25 | ● | ● | ● | ● | ● |  | ● |
| 3 | 8 | ● | ● | ● | ● | ● | ● | ● |
| 4 | 3 | ● | ● | ● | ● | ● | ● |  |
| 5 | 0.91 | ● | ● | ● | ● |  |  | ● |
| 6 | 0.86 | ● | ● | ● |  | ● |  |  |
| 7 | 0.85 | ● | ● | ● | ● |  |  |  |
| 8 | 0.70 | ● | ● | ● | ● |  | ● | ● |
| 9 | 0.67 | ● | ● | ● |  | ● |  | ● |
| 10 | 0.51 | ● | ● | ● | ● |  | ● |  |

**Supplementary Table 7.**
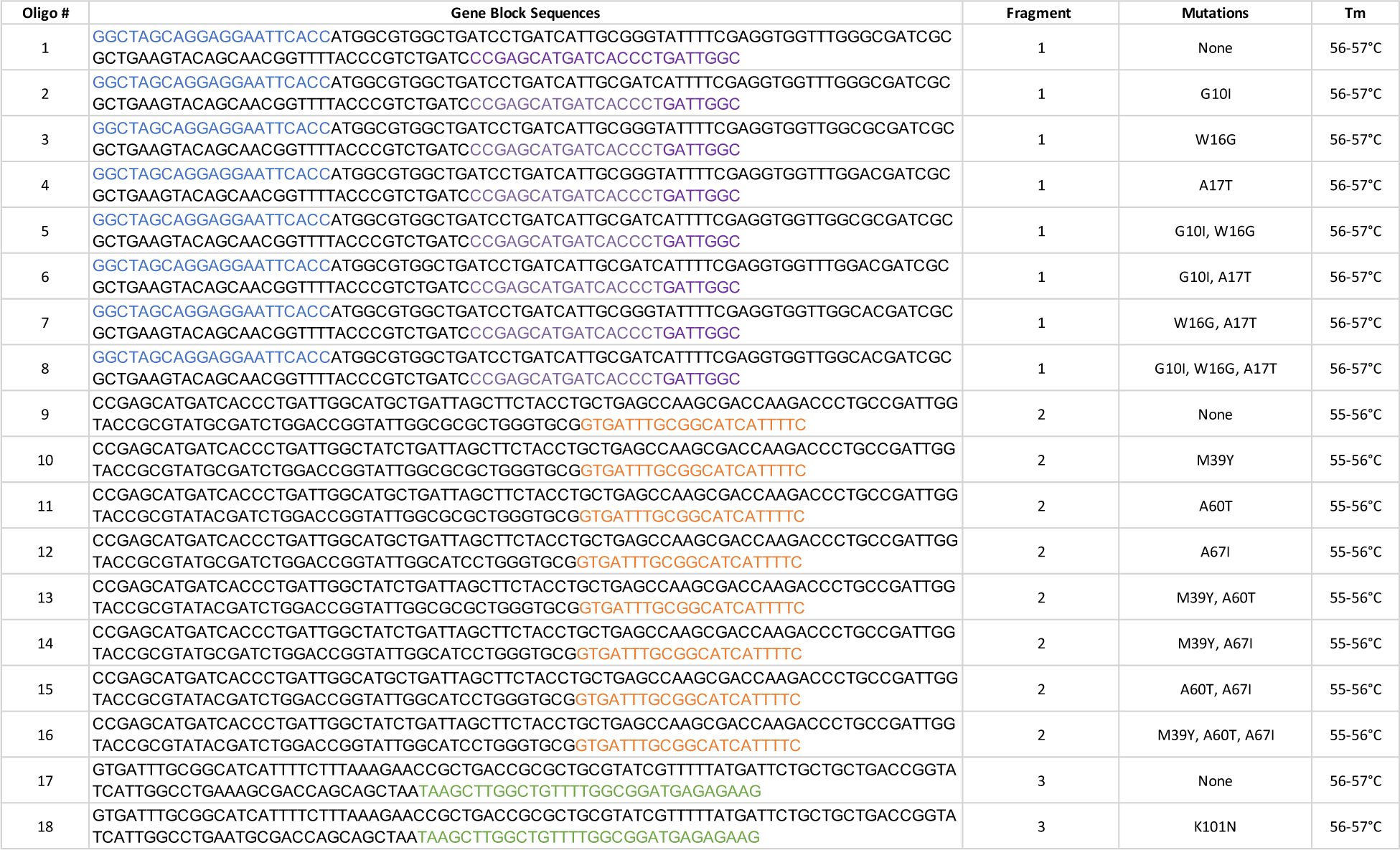
Gene block sequences for combinatorial library assembly.

## Notes

### Competing Interest Statement

The authors have declared no competing interest.

